# γ–Secretase controls the specification of astrocytes from oligodendrocyte precursor cells via Stat3

**DOI:** 10.1101/748772

**Authors:** Jinxing Hou, Huiru Bi, Gang Zou, Zhuoyang Ye, Jing Zhao, Yimin Hu, Jun Lu, Guiquan Chen

**Author notes:** These authors contributed equally to this work (Jinxing Hou, Huiru Bi).

## Abstract

Oligodendrocytes (OLs) and astrocytes play critical roles in a variety of brain functions. OL precursor cells (OPCs) are known to give rise to OLs as well as astrocytes. However, little is known about the mechanism by which OPCs determine their specification choice for OLs versus astrocytes in the central nervous system (CNS). Here we show that genetic inhibition of γ-secretase in OPCs reduces OL differentiation but enhances astrocyte specification. Mechanistic analysis reveals that inhibition of γ-secretase results in decreased levels of Hes1, and that Hes1 down-regulates the expression of signal transducer and activator of transcription3 (Stat3) via binding to specific regions of its promoter. We demonstrate that conditional inactivation of Stat3 in OL lineages restores the number of astrocytes in γ-secretase mutant mice. In summary, this study identifies a key mechanism which controls OPC’s specification choice for OL versus astrocyte during postnatal development. This γ-secretase-dependent machinery may be essential for the CNS to maintain the population balance between OLs and astrocytes.

## Introduction

As major constituents in the CNS, OLs and astrocytes are essential for a variety of brain functions (Rowitch and Kriegstein, 2010). Whereas OLs are myelin-producing cells (Nave and Trapp, 2008), astrocytes play multiple roles including the formation of the brain blood barrier and inflammatory responses (Freeman, 2010; Molofsky et al., 2012; Rowitch and Kriegstein, 2010). Oligogenesis in the CNS has been extensively studied, and several transcriptional factors are critical for this process (Emery, 2010; Jiang et al., 2016; Li and Richardson, 2016). On one hand, it is believed that neural progenitor cells (NPCs) generate astrocytes at the late gestational stage during cortical development (Freeman, 2010; Imayoshi and Kageyama, 2014; Namihira and Nakashima, 2013; Rowitch and Kriegstein, 2010). On the other hand, the following evidence has shown that OPCs give rise to astrocytes as well. First, glial progenitor cells were reported to generate OLs and astrocytes (Kondo and Raff, 2000; Raff et al., 1983). Second, NG2-expressing cells were found to differentiate into both OLs and astrocytes in the CNS (Belachew et al., 2003; Cai et al., 2007; Huang et al., 2014; Zuo et al., 2018). However, little is known about how OPCs decide their specification choice for OLs versus astrocytes during postnatal development.

Notch receptors and amyloid precursor protein (APP) are main substrates of γ-secretase (De Strooper, 2003; Fukumori and Steiner, 2016). The latter is composed of four subunits including presenilin, presenilin enhancer2 (Pen-2), nicastrin (NCT) and anterior pharynx defective 1 (Aph-1) (De Strooper, 2003; Kimberly et al., 2003). Recent cryo-electron microscopy studies have uncovered structural basis for the recognition of γ-secretase to Notch or APP (Yang et al., 2019; Zhou et al., 2019). It has previously been shown that activation of Notch1 inhibits OL differentiation in the developing optic nerve (Wang et al., 1998), and that conditional inactivation of Notch1 enhances OL differentiation in the CNS (Genoud et al., 2002; Zhang et al., 2009) as well as in the peripheral nervous system (PNS) (Woodhoo et al., 2009). As one of upstream proteases for Notch, γ-secretase has been shown to play important roles in embryonic and adult brains (Acx et al., 2017; Cheng et al., 2019; Dries et al., 2016; Hou et al., 2016; Kim and Shen, 2008; Liu et al., 2017; Saura et al., 2004; Shen and Kelleher, 2007; Tabuchi et al., 2009). Surprisingly, it remains unknown how γ-secretase in OPCs regulates OL and astrocyte development in the CNS.

To address the above question, we generated OL lineages specific *Pen-2* conditional knockout (cKO) and *NCT* cKO mice. We find that these mutant animals exhibit decreased number of CC1-positive (+) cells but increased number of GFAP+ cells in the CNS without evident neuron loss. Lineage-tracing analyses reveal that GFAP+ cells are derived from Cre-expressing OPCs *in vivo* and *in vitro*. We demonstrate that deletion of Pen-2 causes decreased levels of Hes1, and that Hes1 inhibits the expression of Stat3 through binding to its promoter. We discover that elevated Stat3 in OPCs enhances astrocyte differentiation via activation of GFAP. Together, this study uncovers molecular mechanisms by which OPCs choose their specification choice in the CNS during postnatal development.

## Results

### Deletion of Pen-2 causes loss of mature OLs in vivo and in vitro

We generated OL lineages specific *Pen-2* cKO (*Pen-2^f/f^;Olig1-Cre*) mice by breeding *Pen-2^f/f^* with *Pen-2^f/+^;Olig1-Cre* (Fig. 1A). To visualize expression pattern of the Cre recombinase, we generated a reporter mouse in which tdTomato is conditionally expressed (Fig. EV1A). This line was simply referred to as *LSL-tdTomato* hereafter and was crossed to the *Olig1-Cre* mouse (Dai et al., 2014; Fu et al., 2009; Lu et al., 2002). Immunohistochemistry (IHC) data showed that tdTomato was co-stained with Olig2 but not Iba1 and NeuN in Cre-expressing cells in the corpus callosum (CC) in the *Olig1-Cre;LSL-tdTomato* mouse at postnatal day 2 (P2) (Fig. EV1B).

**Figure 1.**
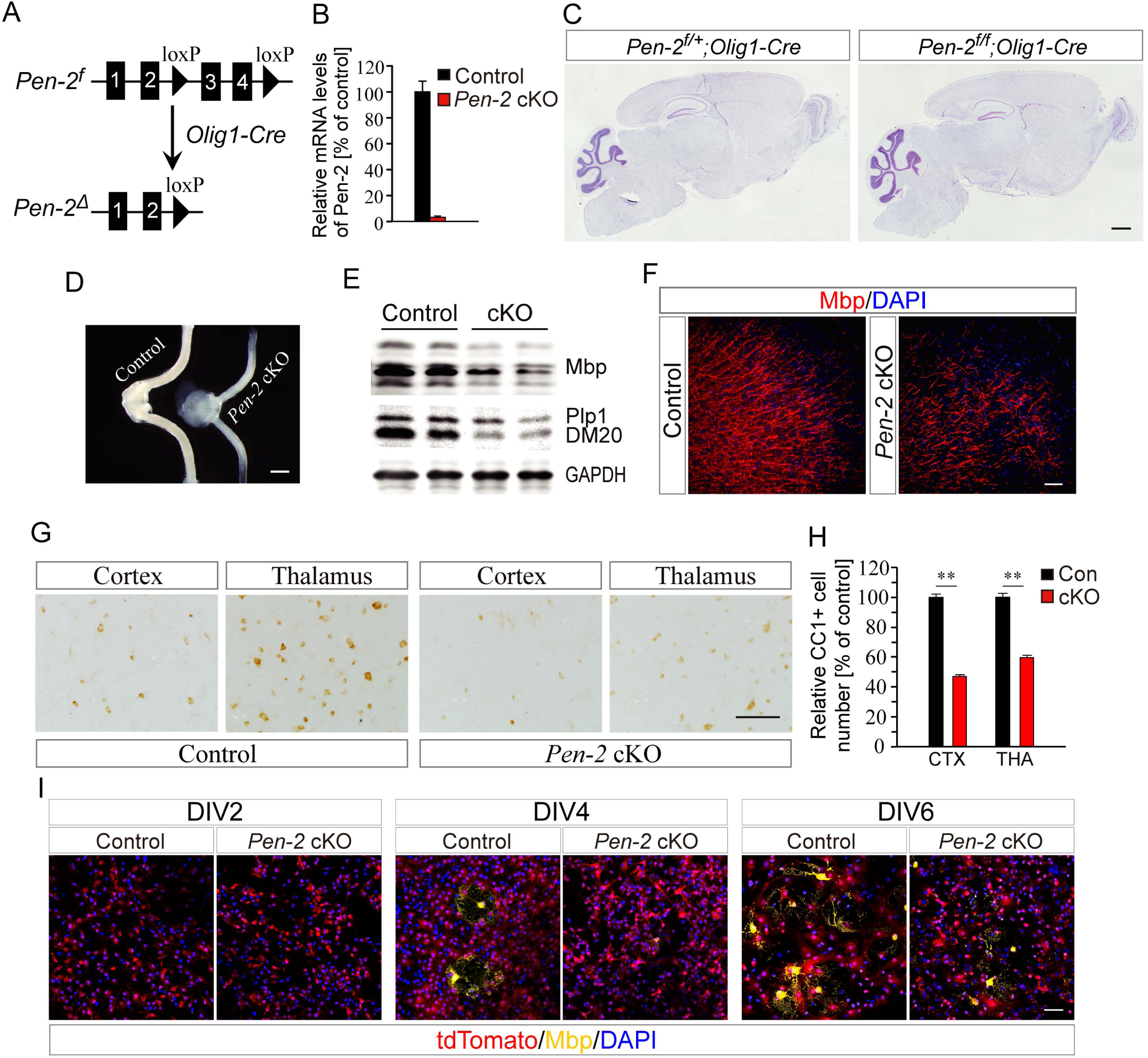
Loss of mature OLs in *Pen-2* cKO mice and in *Pen-2* cKO OL cultures. A. Schematic diagram for the generation of OL lineage specific *Pen-2* cKO mice. B. Quantitative RT-PCR analysis on *Pen-2* mRNA. The FACS technique was used to collect GFP+ cells from *Pen-2^f/+^;Olig1-Cre;mTmG* (control) and *Pen-2^f/f^;Olig1-Cre;mTmG* (*Pen-2* cKO) mice. There was highly significant difference on mRNA levels of *Pen-2* between two groups (Control: n=3; *Pen-2* cKO: n=4; p<0.001). C. Nissl staining. There was no detectable change on brain morphology in *Pen-2* cKO mice at P30 as compared to controls. D. Optic nerves in mice at P21. The optic nerve was translucent in *Pen-2* cKO mice. E. Western analysis on Mbp and Plp1. There were significant differences between control and *Pen-2* cKO mice at P14 (Control: n=3; *Pen-2* cKO: n=4). F. IHC on Mbp. There was qualitative decrease on the immuno-reactivity of Mbp in *Pen-2* cKO mice at P30 as compared to controls. G. IHC on CC1. Brain sections at P30 were used and various brain regions were shown here. H. Relative number of CC1+ cells in 1 mm^2^ area in *Pen-2* cKO mice. There were significant differences on the averaged number of CC1+ cells in the cortex and the thalamus between control and *Pen-2* cKO mice at P30 (Control: n=4; *Pen-2* cKO: n=5; **, p<0.01). I. Co-staining for tdTomato and Mbp using OPC cultures from cortices for *Pen-2^f/+^;Olig1-Cre;LSL-tdTomato* (control) and *Pen-2^f/f^;Olig1-Cre;LSL-tdTomato* (*Pen-2* cKO) mice. There were more Mbp+ cells in *Pen-2* cKO cultures than in controls at DIV4 and DIV6 but not DIV2. Scale bar is 1 mm in (C) and (D), 50 µm in (F) and (G), or 100 µm in (I).

To examine inactivation efficiency of Pen-2, we first obtained *Pen-2^f/+^;Olig1-Cre;mTmG* and *Pen-2^f/f^;Olig1-Cre;mTmG* mice by crossing *mTmG* (Muzumdar et al., 2007) to *Pen-2^f/+^;Olig1-Cre*. RNA samples were prepared using GFP+ cells purified from cortical tissues in the above mice by the fluorescence activated cell sorting (FACS) equipment. Q-PCR analysis revealed very low *Pen-2* mRNA levels in *Pen-2^f/f^;Olig1-Cre;mTmG* (Fig. 1B). Secondly, Western analysis showed that levels for full-length of APP (APP-FL) were not changed but those for C-terminal fragment of APP (APP-CTF) were increased in *Pen-2* cKO (*Pen-2^f/f^;Olig1-Cre*) cortices at P14 and P30 compared with controls (Fig. EV1C). Thus, conditional deletion of Pen-2 in OL lineages significantly inhibited γ-secretase function.

*Pen-2* cKO mice were born in expected Mendelian ratios and survived to adulthood. Nissl staining showed normal brain architecture in *Pen-2* cKO mice at P14 (data not shown) and P30 (Fig. 1C). However, the optic nerve was translucent in *Pen-2* cKO mice (Fig. 1D), suggestive of myelination defect. We then examined myelin-related proteins. First, biochemical analysis revealed reduced levels of Mbp and Plp1 in *Pen-2* cKO cortices at P14 (Fig. 1E) and P30 (data not shown) compared with controls. Second, IHC results showed decreased immuno-reactivity of Mbp (Fig. 1F) and Plp1 (data not shown) in *Pen-2* cKO cortices. Third, relative thickness of Mbp+ fibers was significantly reduced in *Pen-2* cKO mice (Fig. EV1D).

To study whether loss of Pen-2 affected mature OLs, IHC on CC1 was performed. We found that the averaged number of CC1+ cells in the cortex and the thalamus was significantly decreased in *Pen-2* cKO mice at P14 (Fig. EV1E,F: p<0.01) and P30 (Fig. 1G,H: p<0.01) compared with controls, suggesting loss of mature OLs.

We next cultured OPCs using cortices from *Pen-2^f/+^;Olig1-Cre;LSL-tdTomato* (control) and *Pen-2^f/f^;Olig1-Cre;LSL-tdTomato* (*Pen-2* cKO) mice. Western blotting did not detect Pen-2 in *Pen-2* cKO OPC cultures compared with controls (Fig. EV1G). Less Mbp+ cells were observed in *Pen-2* cKO OPC cultures than in controls at DIV4 (differentiation day 4) and DIV6 (Fig. 1I). Overall, the *in vitro* findings were consistent with those from the *in vivo* study.

### Deletion of Pen-2 or NCT results in increased number of OPCs in the CNS

To examine whether OPCs were affected, IHC on Olig2 (Fig. 2A) and Pdgfrα (Fig. 2B) was conducted. We found that the number of Olig2+ cells was increased in the cortex and the thalamus of *Pen-2* cKO mice at P14 compared with controls (Fig. 2C: p<0.01). Moreover, there was significant increase on the total number of Pdgfrα+ cells in the cortex or the thalamus in *Pen-2* cKO mice at P14 (Fig. 2C: p<0.005). Western analysis also showed significantly increased levels of Olig2 and Pdgfrα in P14 *Pen-2* cKO cortices (Fig. 2D: p<0.005).

**Figure 2.**
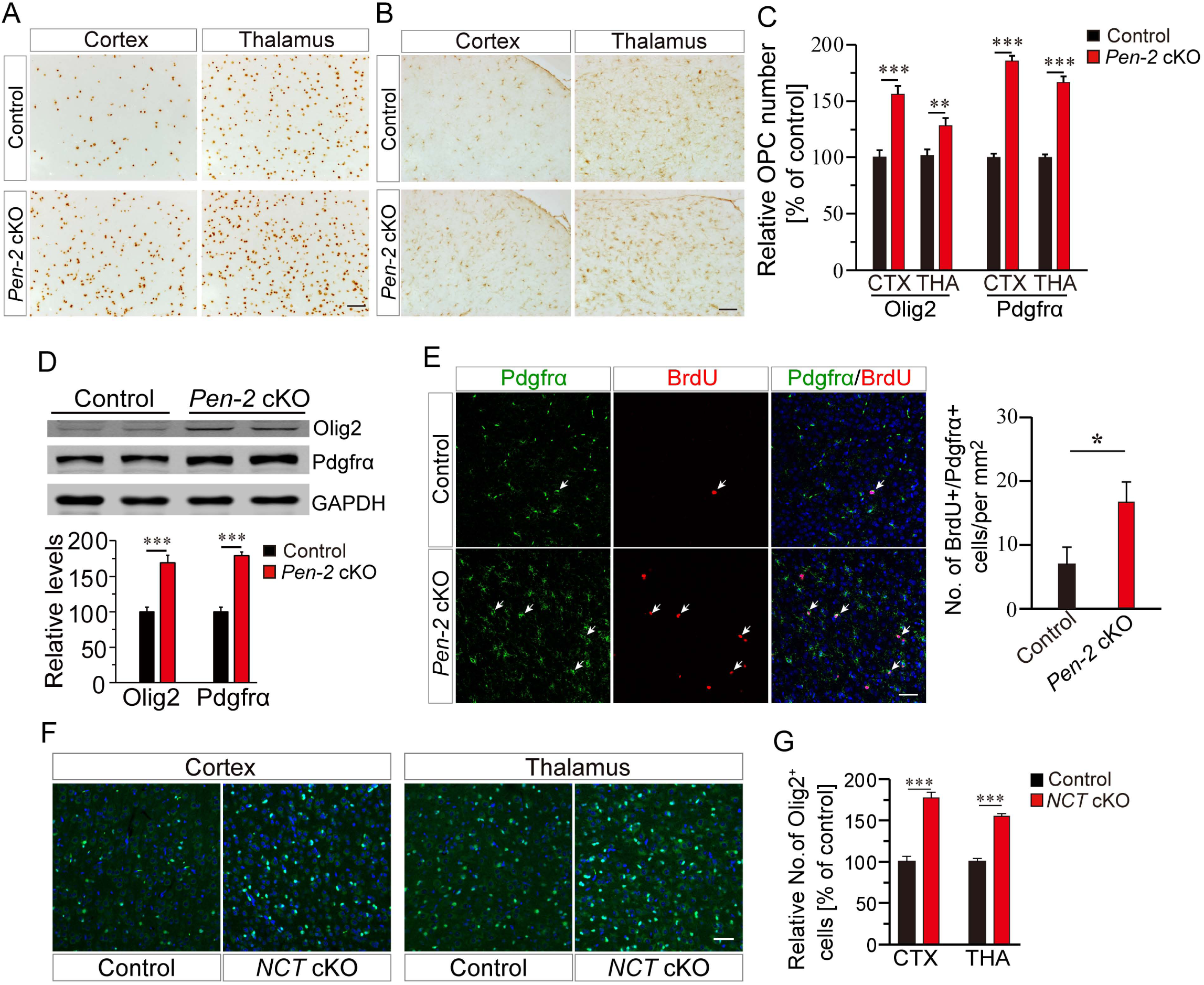
Increased number of OPCs in *Pen-2* cKO and *NCT* cKO mice. (A-B). IHC on Olig2 (A) or Pdgfrα (B) in the cortex and the thalamus for *Pen-2* cKO mice at P14. C. Relative number of Olig2+ and Pdgfrα+ cells. There were significant differences on the averaged number of Olig2+ and Pdgfrα+ cells in 1 mm^2^ area in the cortex and the thalamus between control and *Pen-2* cKO mice at P14 (n=4 mice per group; **, p<0.01; ***, p<0.005). D. Western blotting for Olig2 and Pdgfrα. There were significant differences on levels for Olig2 and Pdgfrα between control and *Pen-2* cKO mice at P14 (Control: n=3; *Pen-2* cKO: n=4; ***, p<0.005). GAPDH served as the loading control. E. Double immuno-staining for Pdgfrα and BrdU. BrdU was injected into P14 mice to label proliferating OPCs. Cells doubly positive for Pdgfrα and BrdU were indicated by arrows. There was a significant difference on the number of Pdgfrα+/BrdU+ cells in the cortex between control and *Pen-2* cKO mice at P14 (Control: n=4; *Pen-2* cKO: n=6; *, p<0.05). F. IHC on Olig2 in the cortex and the thalamus for *NCT* cKO mice at P14. G. Relative number of Olig2+ cells in *NCT* cKO mice. There were significant differences on the averaged number of Olig2+ cells in the cortex and the thalamus between control and *NCT* cKO mice at P14 (n=5 mice per group; ***, p<0.005). Scale bar is 50 µm in (A), (B), (E) and (F).

To find out when the number of OPCs started to get increased in *Pen-2* cKO mice, IHC on Olig2 was carried out using brain sections at P4, P7 and P11 (Fig. EV2A). We observed significantly increased number of Olig2+ cells in the cortex and the thalamus in *Pen-2* cKO mice at P11 but not P4 and P7 (Fig. EV2B). Interestingly, the number of Olig2+ or Pdgfrα+ cells did not differ between controls in *Pen-2* cKO mice at P30 (data not shown). To examine whether proliferation of OPCs was affected, BrdU pulse-labeling was conducted using brains at P14. We observed increased number of cells doubly positive for BrdU and Pdgfrα in *Pen-2* cKO cortices compared with controls (Fig. 2E: p<0.05).

To study whether Pen-2 regulates OPC development in a γ-secretase-dependent manner, we generated OL lineages specific *NCT* cKO mice (Fig. EV2C). We detected decreased levels of NCT (Fig. EV2D) and increased levels of APP-CTF (data not shown) in *NCT* cKO mice by Western blotting. Nissl staining showed normal brain morphology in *NCT* cKO mice compared with controls (Fig. EV2E). Furthermore, Olig2 IHC revealed increased number of Olig2+ cells in *NCT* cKO mice at P14 (Fig. 2F,G: p<0.005). Thus, deletion of NCT caused comparable changes on OPCs to those by loss of Pen-2.

### Deletion of Pen-2 or NCT leads to increased number of astrocytes in the CNS

To examine whether astrocytes were affected, we performed GFAP IHC. First, there was increased immuno-reactivity of GFAP in the cortex and the thalamus of *Pen-2* cKO mice at P14 and P30 compared with controls (Fig. 3A). Second, we observed significantly increased number of GFAP+ cells in the brain of *Pen-2* cKO mice (Fig. 3B: p<0.005). Third, we crossed the *hGFAP-GFP* mouse (Zhuo et al., 1997) to *Pen-2* cKO to obtain *Pen-2^f/f^;Olig1-Cre;GFAP-GFP* (Fig. 3C). The latter exhibited abundant GFP+ cells in the cortex and the thalamus compared with controls (*Pen-2^f/+^;Olig1-Cre;GFAP-GFP*) (Fig. 3D). Fourth, there was increased immuno-reactivity of GFAP in the SC in *Pen-2* cKO mice compared with controls at P14 (Fig. EV3A) and P30 (data not shown).

**Figure 3.**
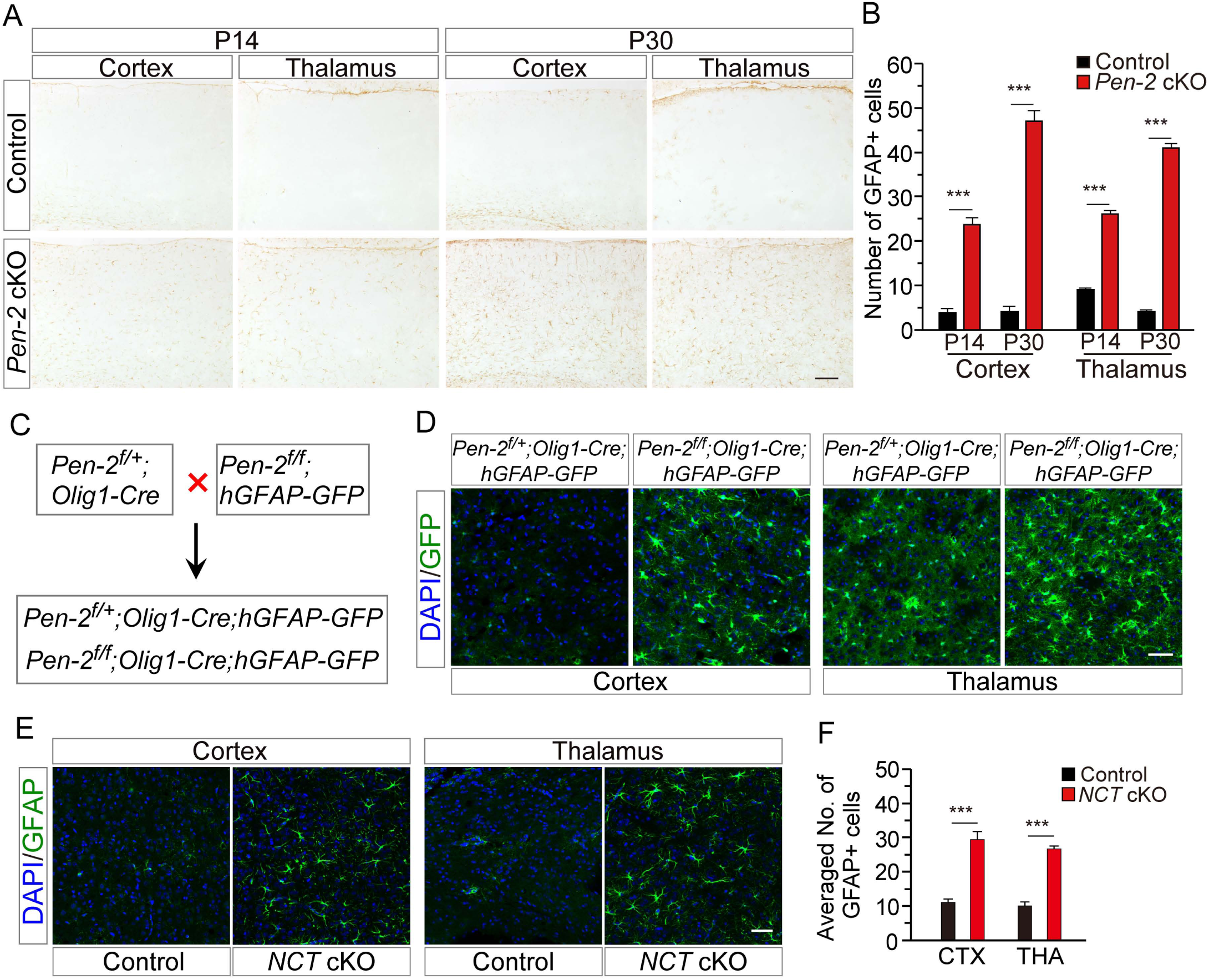
Enhanced astrogenesis in *Pen-2* cKO and *NCT* cKO mice. A. IHC on GFAP in *Pen-2* cKO mice. Brain sections at P14 and P30 were used. There was increased immuno-reactivity of GFAP in the cortex and the thalamus in *Pen-2* cKO mice. B. Averaged number of GFAP+ cells in *Pen-2* cKO mice. There was dramatically increased number of GFAP+ cells in *Pen-2* cKO mice at P14 or P30 as compared to controls (n=6 mice per group; ***, p<0.005). C. Breeding plan for generating *Pen-2* cKO mice expressing GFP in astrocytes. D. *In vivo* labeling of astrocytes by GFP. *Pen-2^f/+^;Olig1-Cre;hGFAP-GFP* (control) and *Pen-2^f/f^;Olig1-Cre;hGFAP-GFP* (*Pen-2* cKO) mice were obtained. Strong GFP expression was seen in the cortex and the thalamus in *Pen-2* cKO mice at P14 as compared to controls. E. IHC on GFAP in *NCT* cKO mice. There was increased immuno-reactivity in the cortex and the thalamus in *NCT* cKO mice at P14 compared with controls. F. Averaged number of GFAP+ cells in *NCT* cKO mice. There was significant increase in the cortex and the thalamus as compared to controls (n=3 mice per group; ***, p<0.005). Scale bar is 100 µm in (A) or 50 µm in (D) and (E).

Western blotting showed increased GFAP levels in cortical samples of *Pen-2* cKO mice at P14 (Fig. EV3B). To examine whether there was abnormal proliferation in astrocytes, BrdU was injected into mice at P14. However, GFAP+/BrdU+ cells were not observed in the brain of control and *Pen-2* cKO mice (Fig. EV3C). Moreover, we found that GFAP+ cells were immuno-positive for GS (glutamine synthetase) and S100β in *Pen-2* cKO mice (Fig. EV3D,E). In addition, we performed GFAP IHC using brain sections of *NCT* cKO mice (Fig. 3E). We found that the averaged number of GFAP+ cells was increased in the cortex and the thalamus in *NCT* cKO mice at P14 (Fig. 3F: p<0.005) and P30 (Fig. EV3F,G: p<0.005).

### Astrocytes are derived from Cre-expressing OPCs in vivo and in vitro

To identify the underlying mechanisms, we first performed lineage-tracing experiments by crossing *Pen-2* cKO to the *LSL*-*tdTomato* reporter mouse (Fig. EV4A). There were abundant tdTomato+ cells but scarce GFAP+ cells, which were negative for tdTomato, in the cortex of *Pen-2^f/+^;Olig1-Cre;LSL-tdTomato* (control) at P14 (Fig. 4A). In contrast, tdTomato+ cells and GFAP+ cells were widely observed in the cortex and the thalamus in *Pen-2^f/f^;Olig1-Cre;LSL-tdTomato* (*Pen-2* cKO) (Fig. 4B). Moreover, we found that GFAP was co-stained with tdTomato in *Pen-2^f/f^;Olig1-Cre; LSL-tdTomato* mice (Fig. 4B,C), indicating that GFAP+ astrocytes are generated from Cre-expressing progenitors. Second, we conducted double staining for Olig2 and GFAP using brain sections at P12. We observed a large quantity of Olig2+/GFAP+ cells in the thalamus (Fig. 4D) and the cortex (Fig. EV4B,C) in *Pen-2^f/f^;Olig1-Cre* mice. Third, we detected many Pdgfrα+/GFAP+ cells in the brain of *Pen-2^f/f^;Olig1-Cre* at P12 (Fig. EV4D). It is likely that Olig2+/GFAP+ or Pdgfrα+/GFAP+ cells represent OPCs which were at the transition stage from OPCs to astrocytes.

**Figure 4.**
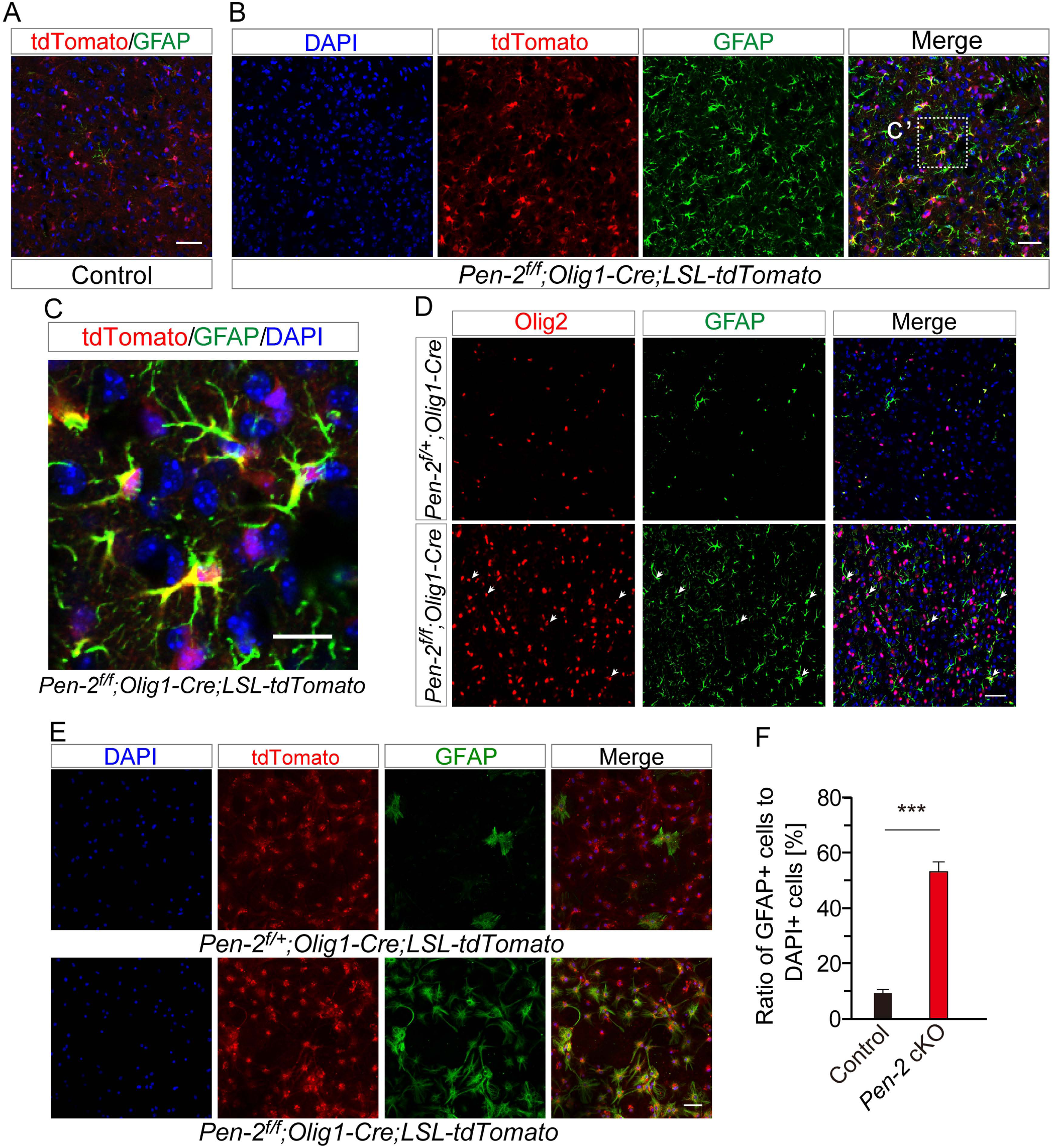
Generation of astrocytes from OPCs in *Pen-2* cKO mice and in *Pen-2* cKO OPC cultures. (A-B) Double staining for GFAP and tdTomato in the cortex of *Pen-2^f/+^;Olig1-Cre; LSL-tdTomato* (control) (A) and *Pen-2^f/f^; Olig1-Cre;LSL-tdTomato* (*Pen-2* cKO) (B) mice at P14. GFAP was co-stained with tdTomato in *Pen-2* cKO but not control mice. C. The boxed area in (B) was enlarged here for co-localization for GFAP and tdTomato in *Pen-2* cKO mice. D. Double staining for Olig2 and GFAP. As indicated by arrows, there were abundant Olig2+/GFAP+ cells in the cortex of *Pen-2* cKO but not control mice at P12. E. Co-staining for tdTomato and GFAP for OPCs cultured from *Pen-2^f/+^;Olig1-Cre; LSL-tdTomato* (control) and *Pen-2^f/f^;Olig1-Cre;LSL-tdTomato* (*Pen-2* cKO) cortices. There was increased immuno-reactivity of GFAP in *Pen-2* cKO OPCs. F. Ratio of GFAP+ cells to DAPI+ cells in OPC cultures. There was significant difference between *Pen-2^f/+^;Olig1-Cre;LSL-tdTomato* and *Pen-2^f/f^;Olig1-Cre;LSL-tdTomato* OPCs at DIV8 (***, p<0.005). Scale bar is 50 µm in (A), (B), (D) and (E) or 10 µm in (C).

To confirm the above *in vivo* findings, we cultured OLs using cortices from *Pen-2^f/+^;Olig1-Cre;LSL-tdTomato* (control) and *Pen-2^f/f^;Olig1-Cre;LSL-tdTomato* (*Pen-2* cKO) mice. Whereas there were scarce cells co-stained with tdTomato and GFAP in control cultures at DIV8, abundant tdTomato+/GFAP+ cells were observed in *Pen-2* cKO cultures (Fig. 4E). Cell counting results showed significantly increased number of GFAP+ cells in *Pen-2* cKO cultures compared with controls (Fig. 4F). These results again indicated that loss of Pen-2 caused increased astrogenesis, and that astrocytes were derived from Cre-expressing cells.

To study whether there was abnormal cell death to induce reactive astrocytes in *Pen-2* cKO mice, we performed TUNEL staining and cleaved caspase3 (CC3) IHC using brain sections at various ages. First, TUNEL+ cells were not observed in the brain of control and *Pen-2* cKO mice at either P11 (Fig. EV5A), P14 (Fig. EV5A) or P30 (data not shown). Second, CC3+ cells were not detected in control and *Pen-2* cKO mice at the above ages as well (data not shown), suggesting no enhanced apoptotic cell death.

We then examined whether there were changes on neurons and microglia in *Pen-2* cKO mice. First, NeuN immuno-reactivity was comparable between control and *Pen-2* cKO mice at P30 (Fig. EV5B). There was no significant difference on the number of NeuN+ cells in the cortex between control and *Pen-2* cKO mice (Fig. EV5B), suggesting no detectable neuron loss in *Pen-2* cKO mice. Western analysis also revealed comparable levels for postsynaptic density protein 95 and synaptophysin between two groups of mice (data not shown). Second, we did not observe significant change on the number of Iba1+ cells in *Pen-2* cKO cortices compared with controls at P14 (Fig. EV5C). Third, the number of NeuN+ cells was not changed in the cortex (Fig. EV5D) or the thalamus (data not shown) in *NCT* cKO mice compared with controls.

### Pen-2 regulates Olig2 expression via Hes1

In line with the finding on Olig2+ cell number (Fig. 2A), *Olig2* mRNA levels were also increased in *Pen-2* cKO cortices at P14 (Fig. EV5E: p<0.01). Since *Pen-2* cKO mice at P11 exhibited increased GFAP immuno-reactivity (Fig. EV5F), we collected GFP+ cells from *Pen-2^f/+^;Olig1-Cre;mTmG* and *Pen-2^f/f^;Olig1-Cre;mTmG* mice at this age using the FACS (Fig. 5A,B). Q-PCR results confirmed decreased *Hes1* mRNA levels but increased *Olig2* mRNA levels in purified *Pen-2* cKO cells compared with controls (Fig. 5C: p<0.01).

**Figure 5.**
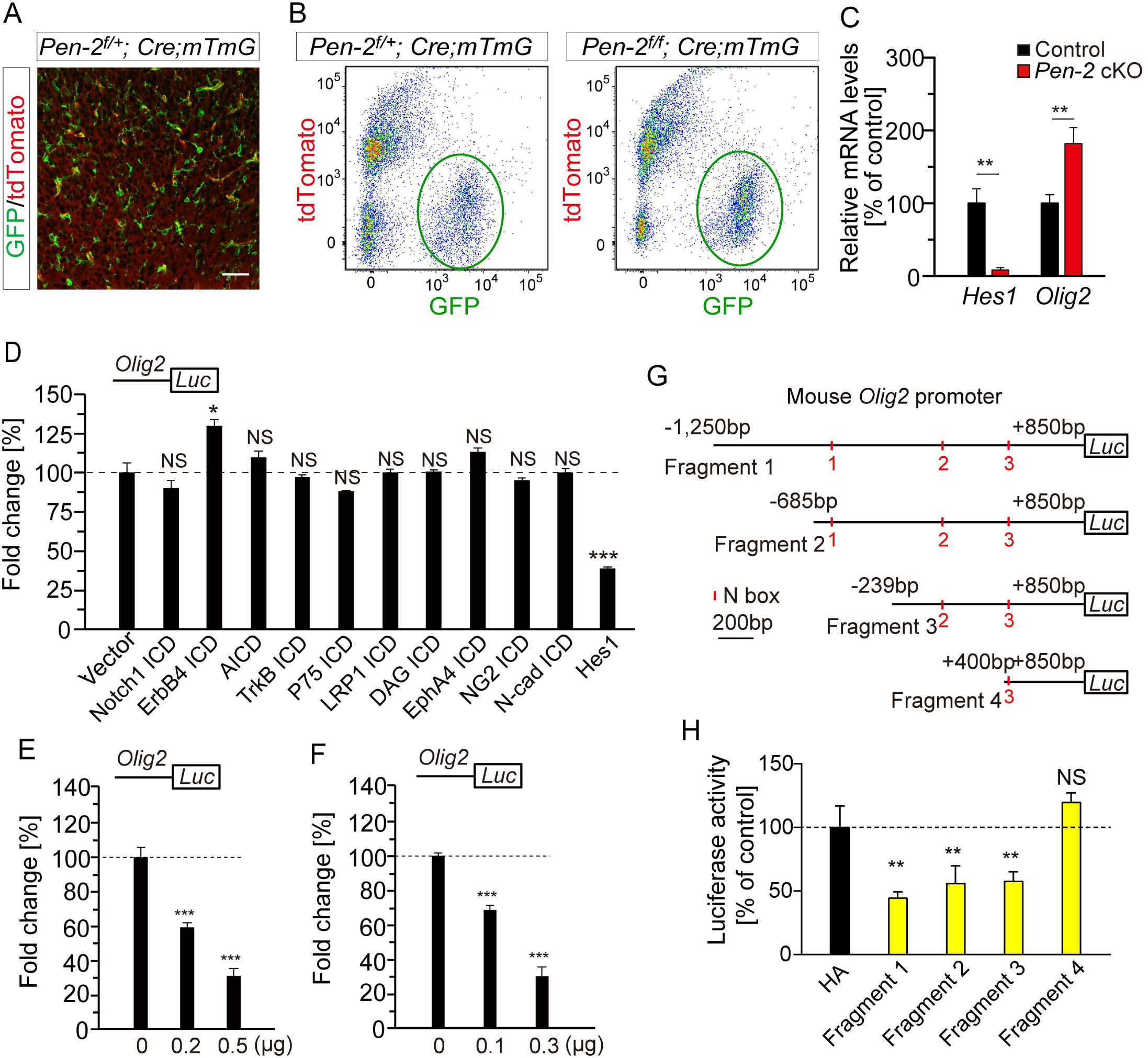
Regulation of Olig2 expression by Hes1. A. Representative images for double staining on GFP and tdTomato in control mice expressing tdTomato and GFP (*Pen-2^f/+^;Olig1-Cre;mTmG*) at P11. Scale bar is 50 µm. B. Purification of *Pen-2* cKO cells expressing GFP by FACS. Cells were collected from *Pen-2^f/+^;Olig1-Cre;mTmG* (control) and *Pen-2^f/f^;Olig1-Cre;mTmG* (*Pen-2* cKO) cortices at P11. C. Q-PCR analysis on *Hes1* and *Olig2* in FACS-purified GFP+ cells. Levels of *Hes1* were decreased but those of *Olig2* were increased in *Pen-2* cKO cells (Control: n=3 mice; *Pen-2* cKO: n=4; **, p<0.01). D. Luciferase assay on *Olig2* promoter activity using HEK293T cells. Expression of Notch1 ICD, APP ICD, TrkB ICD, P75 ICD, LRP1 ICD, DAG ICD, EphA4 ICD, NG2 ICD or N-cadherin ICD did not significantly affect the activity of the *Olig2* promoter (n=4 replicates; p>0.1). Expression of Hes1 significantly inhibited luciferase activity of the *Olig2* promoter (n=4 replicates; p<0.005). (E-F) Dosage-dependent inhibition effect on *Olig2* promoter activity by Hes1 in HEK293T cells (E) or OPC cultures (F) (n=4 replicates; ***, p<0.005). G. Luciferase assay on various fragments of the *Olig2* promoter activity. Fragments 1, 2, 3 and 4 contained the region of −1250bp, −685bp, −239bp or 400bp to 850bp in the *Olig2* promoter, respectively. H. The expression of Hes1 inhibited the luciferase activity of Fragments 1, 2 and 3 but not 4 in HEK293T cells (n=4 replicates; **, p<0.01; NS, not significant).

Notch, APP, ErbB4 and other substrates are processed by γ–secretase to produce different types of intracellular domains (ICDs) (De Strooper, 2003; Lee et al., 2002; Mei and Nave, 2014). To examine whether any of these γ–secretase cleavage products directly regulates Olig2 expression, plasmid expressing Notch1 ICD, AICD, ErbB4 ICD, TrkB ICD, p75 neurotrophin receptor ICD (P75 ICD), LRP1 ICD, DAG ICD, EphA4 ICD, NG2 ICD, N-cadherin ICD or Hes1 was co-transfected with the luciferase reporter under the *Olig2* promoter in HEK293T cells. We found that the *Olig2* promoter activity was not changed by any of these ICDs but was inhibited by Hes1 (Fig. 5D). In addition, we observed a dosage-dependent effect on the *Olig2* promoter activity by Hes1 in HEK293T cells (Fig. 5E). Similar dosage-dependent results were obtained in cultured OPCs as well (Fig. 5F).

To further explore molecular mechanisms, we performed fragment analysis experiments for the *Olig2* promoter (Fig. 5G). Hes1 is a well-known bHLH transcription factor and binds to the N box (CACNAG) but not the E box (CANNTG) in the promoter of its targeted genes (Kageyama et al., 2007; Ohsako et al., 1994). We designed four plasmids containing different regions of the *Olig2* promoter to conduct luciferase experiments. There were three N boxes in Fragments 1 and 2, two in Fragment 3 but none in Fragment 4 (Fig. 5G). We found that expression of Hes1 inhibited luciferase activities for Fragments 1, 2 and 3 but not 4 (Fig. 5H). Thus, the −239bp to +400bp region of the *Olig2* promoter was critical for Hes1-dependent repression on Olig2 expression.

### Pen-2 regulates GFAP expression through Stat3

To conduct molecular analysis for abnormal astrogenesis, we first examined mRNA levels for transcriptional factors important for astrocyte differentiation, Nfia and Stat3, in mice at P11. Q-PCR analyses on RNA samples from FACS-purified cortical GFP+cells revealed increased levels of *Stat3* but not *Nfia* in *Pen-2^f/f^;Olig1-Cre;mTmG* (*Pen-2* cKO) compared with *Pen-2^f/+^;Olig1-Cre;mTmG* (control) mice (Fig. EV6A). To test whether Stat3 was involved *in vivo*, we performed Stat3 IHC. We found that Stat3 immuno-reactivity was increased in the cortex (Fig. 6A) and the thalamus (data not shown) in *Pen-2* cKO mice at P14. In line with this, numerous Stat3+/GFAP+ cells were observed in the brain of *Pen-2* cKO but not control mice (Fig. 6A). Western analysis revealed increased levels of Stat3 in *Pen-2* cKO mice as compared to controls (Fig. 6B). Q-PCR results confirmed significantly increased mRNA levels of *Stat3* and *GFAP* in *Pen-2* cKO cortices at P14 compared with controls (Fig. 6C). There were increased mRNA levels for *NG2* and *Pdgfra* as well (Fig. EV6B: p<0.05). We found that protein levels of Stat3 and GFAP were increased in *Pen-2* cKO OPC cultures compared with controls (Fig. 6D).

**Figure 6.**
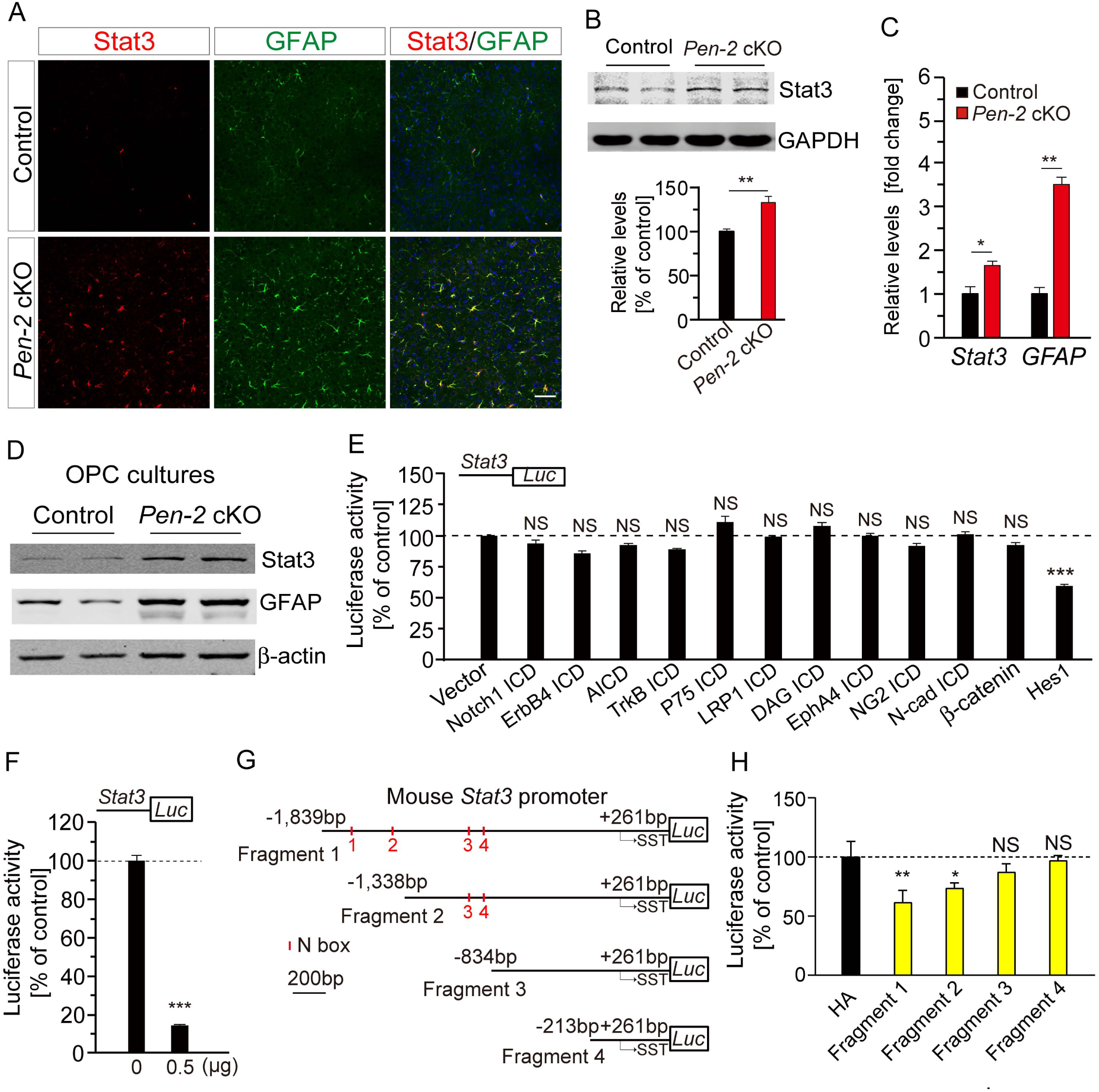
Increased levels of Stat3 and GFAP by Pen-2-dependent regulation on Hes1. A. Double staining for Stat3 and GFAP. The immuno-reactivity of Stat3 was increased in the cortex of *Pen-2* cKO mice at P14. GFAP-expressing cells were immuno-positive for Stat3 in *Pen-2* cKO mice. There were no GFAP+/Stat3+ cells in the cortex of control mice. Scale bar is 50 µm. B. Western analysis on Stat3 in the cortex. There was significant difference between control and *Pen-2* cKO mice at P14 (Control: n=3; *Pen-2* cKO: n=4; **, p<0.01). GAPDH served as the loading control. C. Q-PCR analyses on *Stat3* and *GFAP* mRNAs in the cortex. There were significant differences on their mRNA levels between control and *Pen-2* cKO mice at P14 (n=4 mice per group; *, p<0.05; **, p<0.01). D. Western blotting on Stat3 and GFAP in cultured OPCs. There were increased levels of Stat3 and GFAP in *Pen-2* cKO cell cultures at DIV8 compared with controls (n=4 per group). E. Luciferase assay on *Stat3* promoter activity using HEK293T cells. Expression of Notch1 ICD, APP ICD, TrkB ICD, P75 ICD, LRP1 ICD, DAG ICD, EphA4 ICD, NG2 ICD, N-cad ICD or β-catenin did not significantly affect the activity of the *Stat3* promoter (n=4 replicates; p>0.1). Expression of Hes1 significantly inhibited luciferase activity of the *Stat3* promoter (n=4 replicates; p<0.005). F. Inhibition on *Stat3* promoter activity by Hes1 in OPC cultures (n=4 replicates; ***, p<0.005). G. Luciferase assay on various fragments of the *Stat3* promoter activity. Fragments 1, 2, 3 and 4 contained the region of −1839bp, −1338bp, −834bp or −213bp to 261bp in the *Stat3* promoter, respectively. H. The expression of Hes1 inhibited the luciferase activity of Fragments 1 and 2 but not 3 and 4 in HEK293T cells (n=4 replicates; *, p<0.05; **, p<0.01; NS, not significant).

Moreover, we observed increased Stat3 protein levels in cortical samples of P14 *NCT* cKO mice compared with controls (Fig. EV6C: p<0.01). Levels of GFAP, Olig2 and Pdgfrα but not NeuN were increased in *NCT* cKO mice (Fig. EV6D, p<0.01). Overall, molecular changes in *NCT* cKO mice were comparable to those in *Pen-2* cKOs.

To investigate whether Pen-2 may regulate Stat3 expression via one of the γ–secretase cleavage products, we conducted a number of luciferase experiments using HEK293T cells. However, neither ICD nor β-catenin inhibited the *Stat3* promoter activity (Fig. 6E). In contrast, expression of Hes1 significantly reduced the *Stat3* promoter activity (Fig. 6E), and there was a dosage-dependent effect by Hes1 in HEK293T cells (data not shown). Moreover, we found that expression of Hes1 decreased *Stat3* promoter activity in cultured OPCs as well (Fig. 6F). To further analyze the underlying molecular mechanisms, we generated four plasmids containing different fragments of the *Stat3* promoter. Four N-boxes were included in Fragment 1, two in Fragment 2 but none in Fragments 3 and 4 (Fig. 6G). We found that luciferase activities were inhibited for Fragments 1 and 2 (Fig. 6H). Therefore, the −1338bp to −834bp region of the *Stat3* promoter was critical for Hes1 regulation.

To find out whether GFAP expression was regulated by any γ–secretase cleavage product, HEK293T cells were co-transfected with plasmids expressing luciferase system driven by the *GFAP* promoter and one of the above ICDs. However, neither ICD nor Hes1 significantly inhibited the promoter activity of *GFAP* (Fig. EV6E). In contrast, expression of activated Stat3, human Stat3 or mouse Stat3 but not inactive Stat3 increased the promoter activity of *GFAP* (Fig. EV6F).

### Conditional deletion of Stat3 restores astrocyte population in Pen-2 cKO mice

To verify the importance of the STAT3 signaling in Pen-2-dependent astrogenesis, we conducted a rescue experiment by crossing *Pen-2* cKO to *Stat3^f/f^*. We generated *Pen-2/Stat3* double conditional KO (cDKO) mice in which Pen-2 and Stat3 were both inactivated in OL lineages (Fig. EV7A). As expected, Stat3 levels were decreased in the cortex of *Stat3* cKO and *Pen-2/Stat3* cDKO mice at P14 (Fig. EV7B,C). We found that deletion of Pen-2, Stat3 or Pen-2/Stat3 together did not affect protein levels of APP (Fig. EV7B,C). In contrast, APP-CTFs were accumulated in the cortex of *Pen-2* cKO and *Pen-2/Stat3* cDKO mice (Fig. EV7B,C). Nissl staining revealed no detectable change on brain morphology in either *Stat3* cKO or *Pen-2/Stat3* cDKO mice at P14 compared with their littermate controls (Fig. 7A). Thus, deletion of Stat3 in OPCs did not significantly affect brain development.

**Figure 7.**
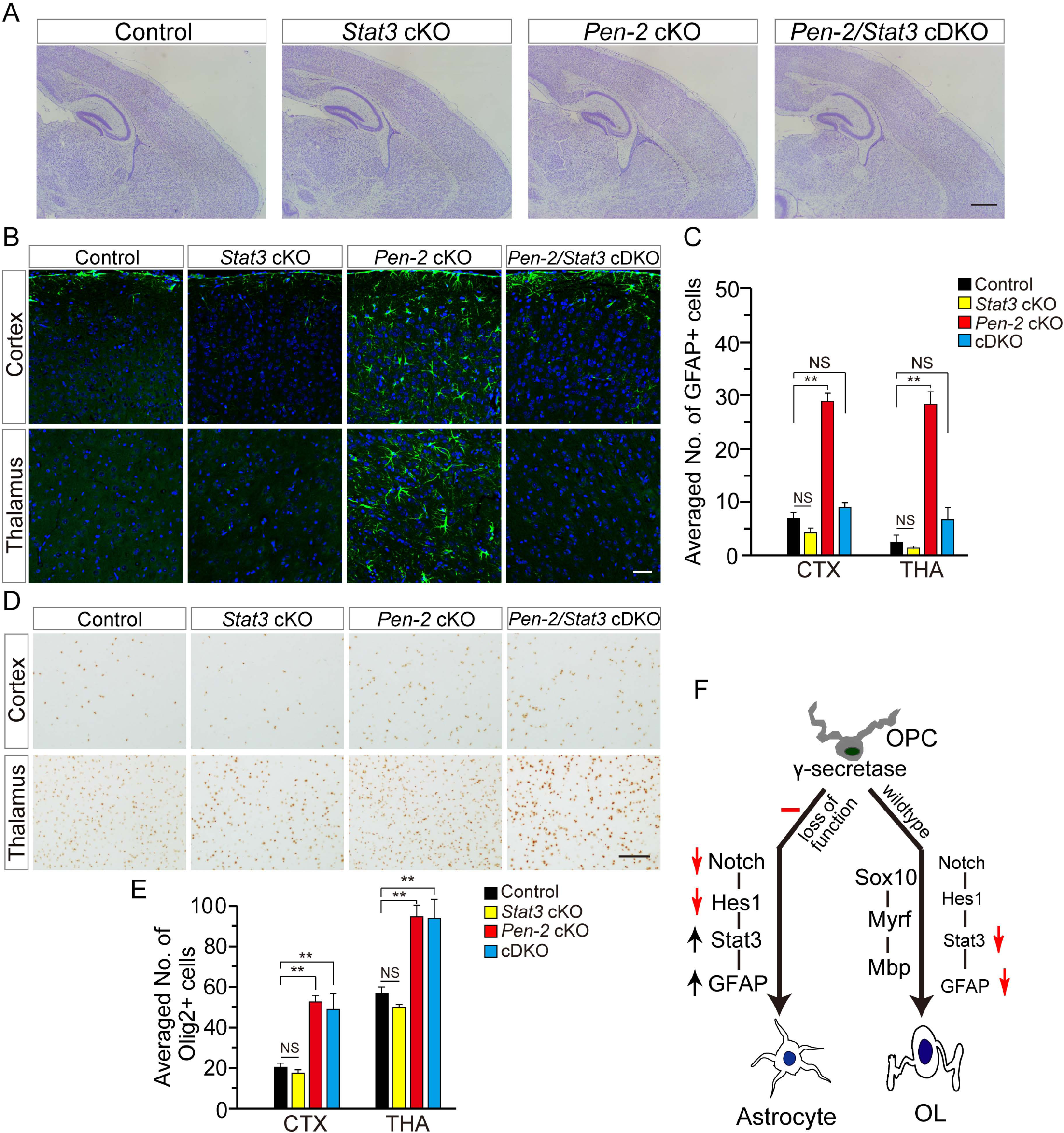
Restoration of astrocyte population in *Pen-2* cKO mice through OL lineages specific deletion of Stat3. A. Nissl staining on brain sections for control, *Stat3* cKO, *Pen-2* cKO and *Pen-2/Stat3* cDKO mice. There were no detectable changes in *Pen-2/Stat3* cDKO cortices as compared to other groups. B. IHC on GFAP. Brain sections for control, *Stat3* cKO, *Pen-2* cKO and *Pen-2/Stat3* cDKO mice at P14 were used. The immuno-reactivity of GFAP was diminished in the cortex of *Pen-2/Stat3* cDKO mice as compared to *Pen-2* cKOs. C. Averaged number of GFAP+ cells in the cortex and the thalamus for mice at P14. There was significant difference between *Pen-2* cKO and *Pen-2/Stat3* cDKO mice (Control: n=6; *Stat3* cKO: n=3; *Pen-2* cKO: n=5; *Pen-2/Stat3* cDKO: n=3; **, p<0.01; NS, not significant). D. IHC on Olig2 in the cortex and in the thalamus. The immuno-reactivity of Olig2 was comparable between *Pen-2* cKO and *Pen-2/Stat3* cDKO mice. E. Averaged number of Olig2+ cells in the cortex and the thalamus. There was no significant difference on the number of Olig2+ cells between *Pen-2* cKO and *Pen-2/Stat3* cDKO mice (**, p<0.01; NS, not significant). F. A cellular model depicts the relevant mechanism to regulate the specification of astrocytes from OPCs. Hes1-dependent repression on Stat3 expression is central to this mechanism. See the Discussion section for detailed description. Scale bar is 500 µm in (A) or 50 µm in (B) and (D).

GFAP IHC showed that the averaged number of GFAP+ cells was remarkably decreased in the cortex and the thalamus of *Pen-2/Stat3* cDKO mice as compared to *Pen-2* cKOs (Fig. 7B,C: p<0.01). In line with this finding, protein levels of GFAP were significantly reduced in *Pen-2/Stat3* cDKO mice compared with *Pen-2* cKOs (Fig. EV7B,C: p<0.01). Therefore, abnormal astrogenesis in *Pen-2* cKO mice was prevented by deletion of Stat3. Western blotting showed that levels of Olig2 were not reduced in *Pen-2/Stat3* cDKO mice compared with *Pen-2* cKOs (Fig. EV7B,C). In addition, the averaged number of Olig2+ cells was not reduced in *Pen-2/Stat3* cDKO mice compared with *Pen-2* cKOs (Fig. 7D,E), suggesting that deletion of Stat3 did not affect OPCs.

## Discussion

Whereas OLs are generated from OPCs, astrocytes are from NPCs as well as OPCs (Belachew et al., 2003; Cai et al., 2007; Freeman, 2010; Huang et al., 2014; Nagao et al., 2016; Zhang et al., 2016; Zuo et al., 2018). Although previous evidence has revealed a negative regulation of OL differentiation by Notch1 (Genoud et al., 2002; Wang et al., 1998; Woodhoo et al., 2009; Zhang et al., 2009), it remains unknown whether γ-secretase plays the same role as Notch1 does in this process. Here we show that deletion of Pen-2 or NCT in OPCs results in loss of mature OLs but enhanced astrogenesis in the CNS (Figs.1-4). We report that deletion of Pen-2 inhibits the Notch signaling but activates Stat3 *in vitro* and *in vivo* (Figs.5-6). We demonstrate that Stat3 is a key mediator for astrocyte specification caused by loss of Pen-2 (Fig. 7). Therefore, the role of γ-secretase in OL development is different from that of Notch1. This discrepancy may get solved by the following explanations. First, since γ-secretase cleaves all Notch receptors, phenotypes in *Pen-2* cKO and *NCT* cKO mice could be caused by complete disruption of Notch function. Second, the remaining Notch receptors, e.g. Notch2/3/4, may produce partial compensatory effects on astrocyte development in *Notch1* cKO mice (Genoud et al., 2002; Wang et al., 1998; Zhang et al., 2009). In agreement with the OL loss shown in *Pen-2* cKO and *NCT* cKO mice in this study, hypomyelination is reported in a different line of OL lineages specific *NCT* cKO mice (Dries et al., 2016).

It has been shown that OPCs possess multi-potency for differentiation (Belachew et al., 2003; Cai et al., 2007; Huang et al., 2014; Zuo et al., 2018). This study provides further support to this concept. Although mechanisms by which NPCs differentiate into astrocytes have been intensively investigated (Freeman, 2010; Hirai et al., 2012; Imayoshi and Kageyama, 2014), it is not well understood how OPCs choose their differentiation route for OLs or astrocytes in the CNS. Findings in this study suggest that γ-secretase may serve as a key “switch” for OPCs to decide a specification choice. Molecular mechanisms underlying abnormal astrogenesis in γ-secretase mutant mice are elaborated as follows (Fig. 7F). First, deletion of Pen-2 or NCT blocks the cleavage of Notch receptors, which leads to decreased Hes1 levels. Second, Hes1 negatively regulates expression of Stat3 and Olig2 through binding to specific regions of their promoters. Third, elevated Stat3 activates GFAP expression, which triggers OPCs differentiating into astrocytes but not OLs. Fourth, in wildtype OPCs, normal γ-secretase activity promotes Hes1 activation which inhibits the expression of Stat3 and GFAP. Meanwhile, OL-related transcriptional factors activate the Myrf/Mbp signaling to trigger OL differentiation and promote oligogenesis (Fig. 7F).

The γ-secretase-dependent regulation on OPC’s specification fate may have significant physiological impacts. According to findings in this study, γ-secretase inhibits astrocyte specification from OPCs in the CNS under normal physiological condition. Therefore, this mechanism could prevent abnormal astrogenesis and maintain the population balance between OLs and astrocytes in the CNS. Overall, this study together with previous ones (Ge et al., 2002; Wang et al., 1998; Watkins et al., 2008; Woodhoo et al., 2009; Zhang et al., 2009) may advance our understanding on the complexity of the γ-secretase/Notch signaling in the brain. Moreover, since mutations on γ-secretase subunits cause various brain disorders (Dermaut et al., 2002; Forzano et al., 2012; Gana et al., 2012; Sala Frigerio et al., 2005; Shen and Kelleher, 2007; Zhong et al., 2009), findings in this study may provide insights on the developmental etiology for abnormal astrogenesis under these conditions.

## Materials and methods

### Animals

The generation of *Pen-2^f/f^* mice was reported previously (Cheng et al., 2019). To generate *Pen-2* cKO mice (*Pen-2^f/f^;Olig1*-*Cre*), we bred *Olig1*-*Cre* mutant (Lu et al., 2002) with *Pen-2^f/f^* to obtain *Pen-2^f/+^*;*Olig1*-*Cre*. The latter were then crossed to *Pen-2^f/f^* to get *Pen-2* cKO. *Pen-2^f/+^*;*Olig1*-*Cre* served as littermate controls. Similar strategies were used to generate *NCT* cKO (*NCT^f/f^*;*Olig1*-*Cre*) mice. The *LSL-tdTomato* reporter mouse was generated using a strategy shown in Fig. EV1. The *mTmG* mouse (Muzumdar et al., 2007) was used to generate *Pen-2^f/+^;Olig1-Cre;mTmG* and *Pen-2^f/f^;Olig1-Cre;mTmG*. The *hGFAP-GFP* mouse (Zhuo et al., 1997) was used to generate *Pen-2^f/f^;Olig1-Cre;hGFAP-GFP* mice. *Stat3^f/f^* mice were reported previously (Moh et al., 2007). *Pen-2/Stat3* cDKO mice were generated using a breeding strategy shown in Fig. EV7.

The genetic background of the mice was C57BL/6. The animals were group-housed (4-5 per cage) throughout the experimental period and had ad libitum access to food and water. The mice were maintained in an SPF-leveled animal room in the core facility of the Model Animal Research Center (MARC) at Nanjing University. The light-cycle of the animal room was automatically controlled. The animal room was maintained under constant humidity and temperature (25±1℃). Mouse breeding was conducted under IACUC approved protocols at the MARC. All the experiments were performed in accordance with the Guide for the Care and Use of Laboratory Animals of the MARC at Nanjing University.

### Nissl staining

The mice were euthanized with CO_2_, perfused with PBS, fixed in 4% paraformaldehyde overnight at 4℃. Sections were deparaffinized, ethanol rehydrated and treated with 0.1% cresyl-violet for 1 min followed by rinsing with water. Sections were sealed using neutral resin.

### Immunohistochemistry (IHC)

IHC was conducted using a method described previously (Ho et al., 2006). Primary antibodies were listed in Table EV1. For fluorescence IHC, images were captured and analyzed using a Zeiss LSM880 confocal microscope.

### Cell counting

Counting on CC1+, Olig2+, Pdgfrα+, NeuN+ or Iba1+ cells was conducted using 3 brain sections per mouse spaced 400 µm apart. For each section, 2 microscopic fields (20× objective lens of an Olympus BX53 microscope) were randomly selected for the cortex or the thalamus. Images for each field (438.6µm × 330.2µm) were captured and the total number of cells was counted. The serial numbers were averaged across sections and the mean was presented as the averaged cell number in each microscopic field for each animal.

### Brain lysate preparation

Mice were sacrificed at P14 and P30. Cortical samples were prepared using a method described previously (Peng et al., 2010; Peng et al., 2007). Samples were stored at −80℃ until use.

### Western blotting

We used a method described previously (Liu et al., 2017; Tabuchi et al., 2009). Briefly, normalized volumes for cortical protein samples were loaded onto 8%-15% polyacrylamide gels and separated by electrophoresis for about 2 hrs. The gels were transferred into nitrocellulose membrane at 25 volts for 2.5 hrs. Primary antibodies used in this study were included in Table EV1.

### Cell culture and transfection

HEK293T cells were maintained in DMEM (Gibco) containing 10% fetal bovine serum (Lonsera). The cells were transfected using polyethyleneimine. The medium was replaced 6 hrs after transfection and the cells were harvested at 48 hrs after transfection.

### Purification of Pen-2 KO cells by FACS

The cortex was dissected from control (*Pen-2^f/+^;Olig1-Cre;mTmG*) and *Pen2* cKO (*Pen-2^f/f^;Olig1-Cre;mTmG*) mice. Cortical samples were digested by trypsin and became single cell suspension containing Cre+ cells expressing green fluorescence protein and Cre-cells expressing red fluorescence protein. Cell suspensions were used by FACS (BD FACS AriaⅢ) to purify Cre+ cells. The sorting efficiency was larger than 90% for each sample. The resultant cell suspensions were centrifuged (3000 rpm) to discard the supernatants. Cells were treated using TRIzol reagent (Invitrogen) for RNA purification.

### OPC and OL cultures

We used a method described previously (Parras et al., 2004). Briefly, cells from cortices from P8 mice were plated in neurosphere medium (DMEM/F12 containing 2mM L-glutamine, 1×B27, 1×N2, 5 µM HEPES, 0.01% Heparin, 100 µg/ml Penicillin, 0.1mg/ml Streptomycin) supplemented with 20 ng/ml EGF and 20 ng/ml FGFb. Primary neurospheres were cultured for 3-5 days. Next, the culture medium was changed to DMEM/F12 supplemented with 20 ng/ml PDGF-AA and 10 ng/m FGFb to induce OPC differentiation. The medium was changed once every 4 days. 12 days after the OPC culture, 10% FBS and 15nM T3 were added into the culture medium to induce OL differentiation. OLs were further cultured for 4-8 days.

### RNA isolation

Total RNA was isolated from control and *Pen-2* cKO cortices using TRIzol reagent (Invitrogen). The method was described previously (Cheng et al., 2019).

### Quantitative RT–PCR (q-PCR)

Total RNA (1 µg) was reverse transcribed using PrimeScript RT reagent Kit (Takara). Real-time PCR was performed using the ABI StepOne Plus system. Primers used for q-PCR were listed in Table EV2.

### TUNEL staining

Brain sections were blocked with 5% goat serum for 30 min and then treated with the TUNEL BrightGreen Apoptosis Detection Kit (Vazyme, Nanjing, China) at 37 °C for 1hr (Cheng et al., 2014; Tabuchi et al., 2009).

### BrdU pulse-labeling

BrdU (Sigma-Aldrich, B5002) was administered at the concentration of 100 mg/kg. To label proliferating OPCs, BrdU was intraperitoneally injected to mice at P14. Brains were collected 1 hr after the injection.

### Plasmids

Mouse *Hes1*, *Hes5*, *ErbB4 ICD*, *AICD*, *TrkB ICD*, *P75 ICD*, *LRP1 ICD*, *DAG ICD*, *EphA4 ICD*, *DAG ICD*, *NG2 ICD*, *N-cadherin ICD*, *Stat3*, *Olig2* were amplified by PCR from cDNA libraries prepared from mouse brain. *NICD* was purchased from addgene (#26891). These cDNAs were subcloned into pCDNA5-HA plasmids. WT human *Stat3* (*Stat3*) and inactive human *Stat3* (*Stat3*) constructs were kindly provided by Prof. Xinyuan Fu. Primers used here were listed in Table EV2.

### Luciferase assay

The mouse *Stat3* promoter (from −1839bp to +261bp) and the *Olig2* promoter (from −1250bp to +850bp) were cloned from mouse NPCs. The human *GFAP* promoter (from −1702 bp to −5 bp) was cloned from DNAs of the *hGFAP-Cre* mouse. These constructs were inserted into the pGL3-luciferase vector (Promega). *Hes1* was co-transfected with *Stat3*-*Luc, Olig2*-*Luc* or *hGFAP*-*Luc* into HEK293T cells. Lipofectamine 2000 (Life Technologies) was used for transfection. The cells were cultured for 48 hrs and cell extracts were assayed for luciferase activity by the Dual-luciferase Reporter Assay System (Promega). Relative luciferase activity was normalized by the Renilla luciferase as the control.

### Statistical analyses

Data were presented as the mean±SEM. Two-tailed student’s t-test was performed to examine the difference between control and mutant mice. P<0.05 (*) was considered statistically significant.

## Acknowledgments

This work was supported by grants from the National Natural Science Foundation of China (91849113 and 31271123), the Jiangsu Provincial Key Medical Discipline (ZDXKA2016020), the Priority Academic Program Development of Jiangsu Higher Education Institutions (PAPD) and the 2016 “333 Project” Award of Jiangsu Province.

## Author contributions

GC, JL and YH designed experiments. JH, HB, GZ and ZY conducted experiments. JZ provided reagents. JH and HB analyzed the data. GC, JL and YH wrote the manuscript.

## Conflict of interests

The authors declare no competing interests.

---

**Table EV1:**
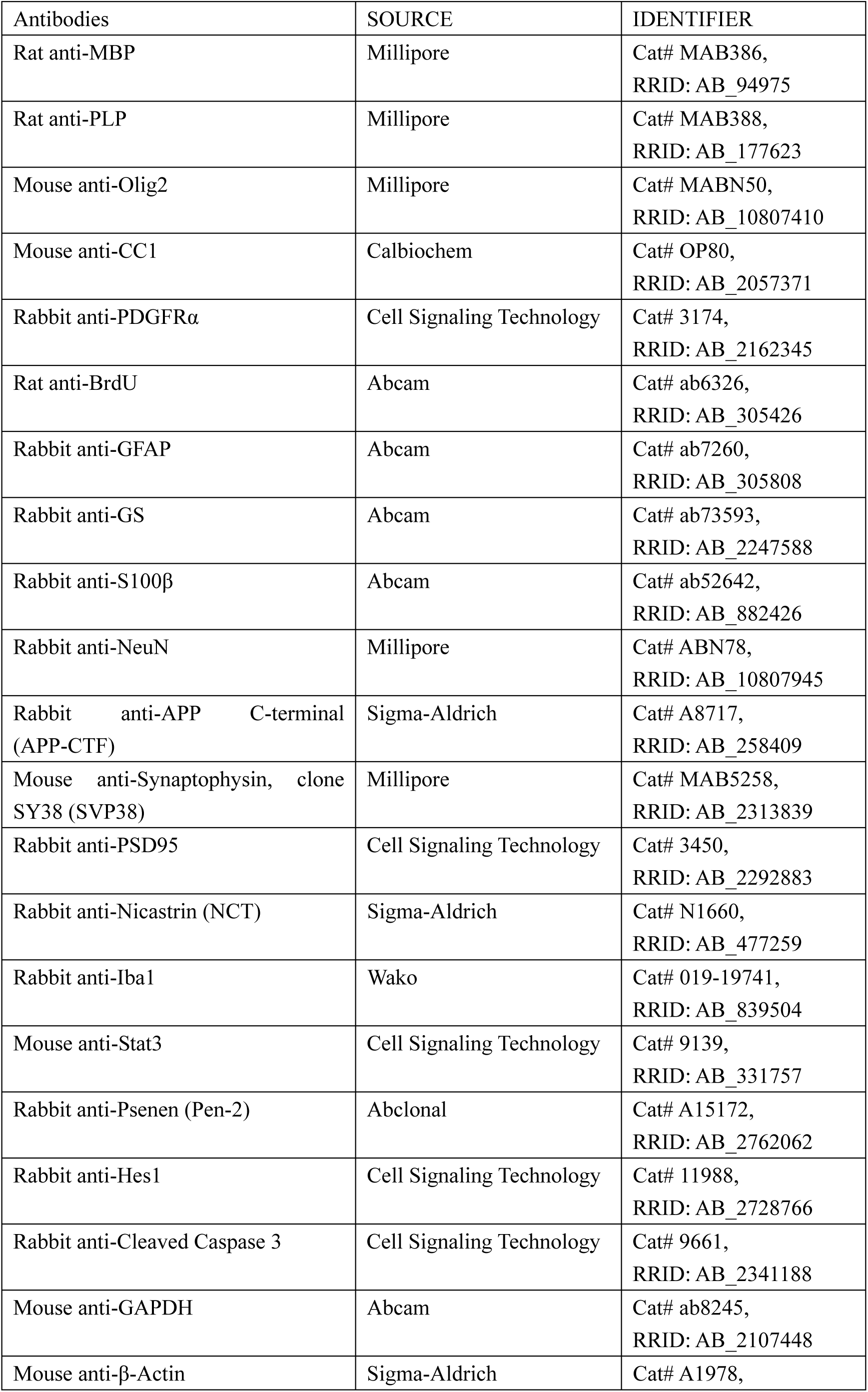
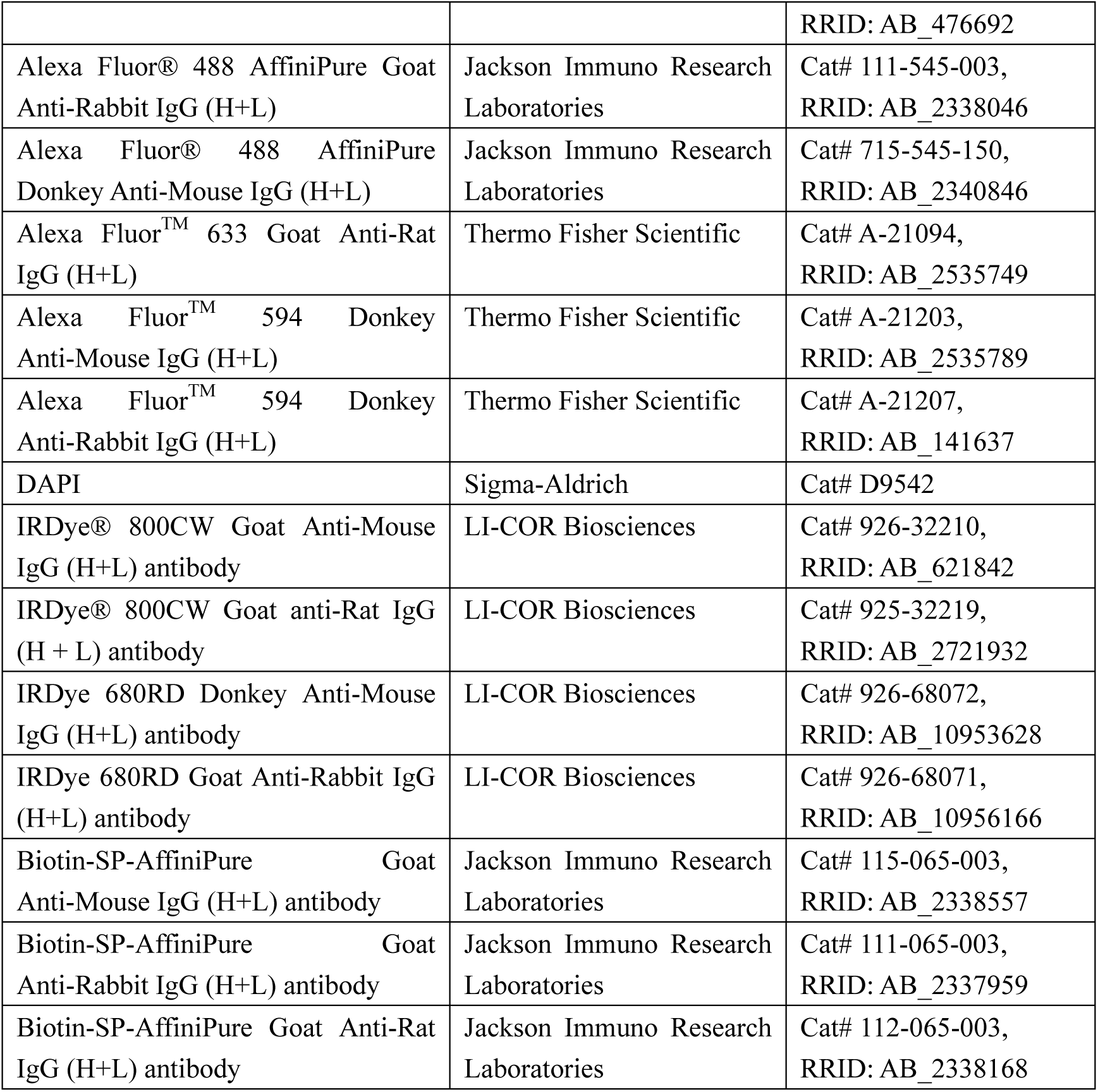
Antibody list.

**Table EV2:**
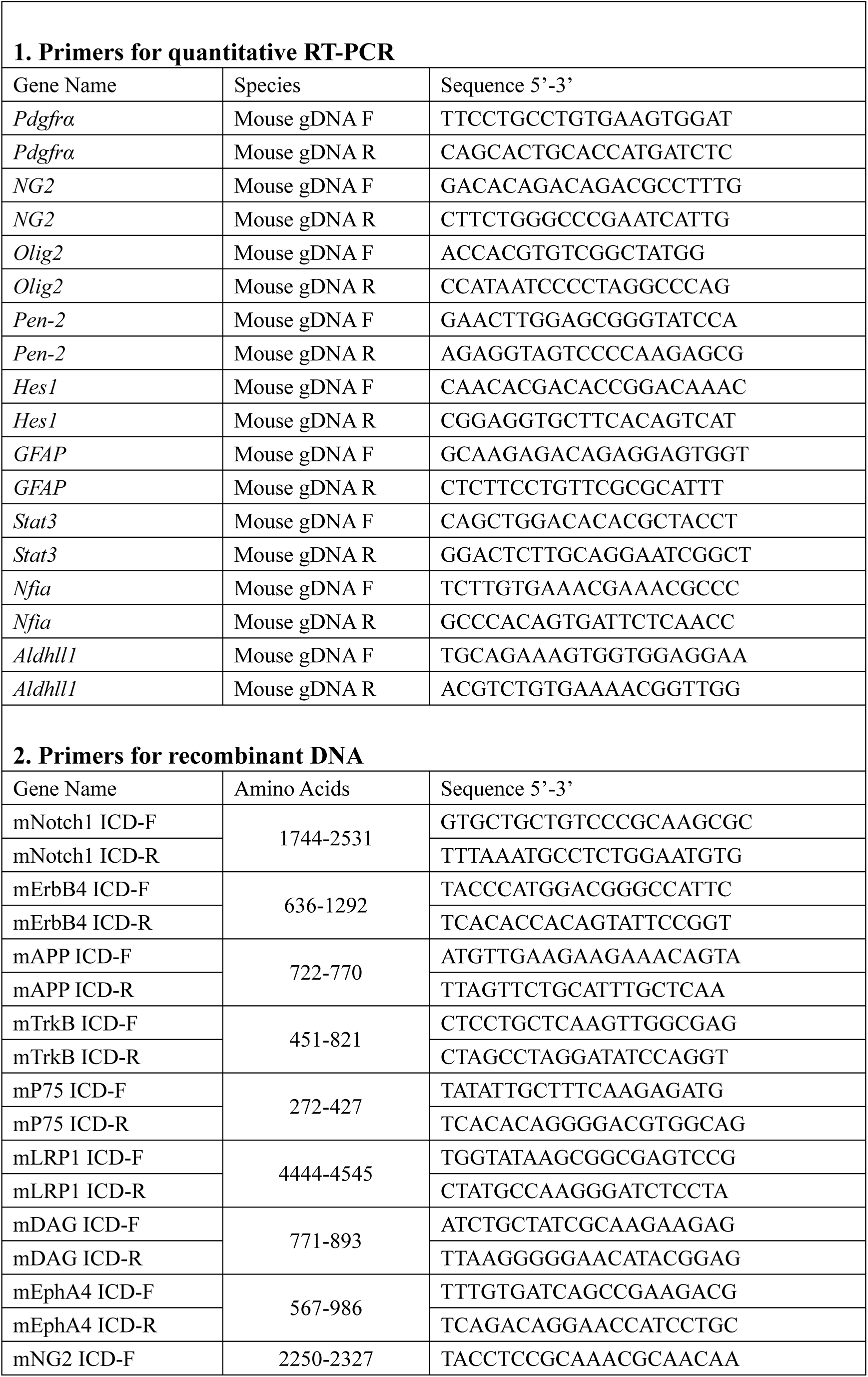
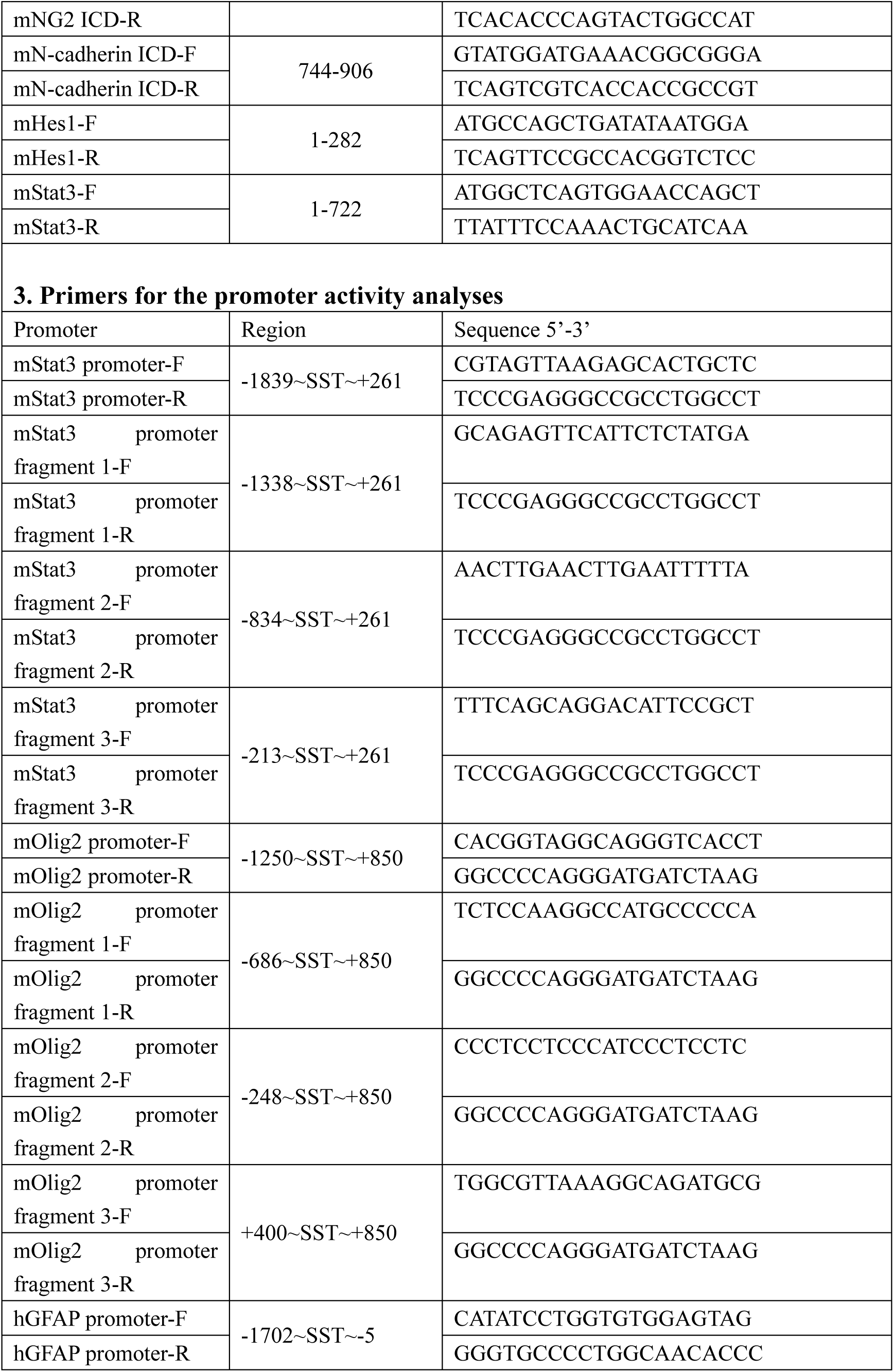
List for primers used in this study.

**Figure EV1.**
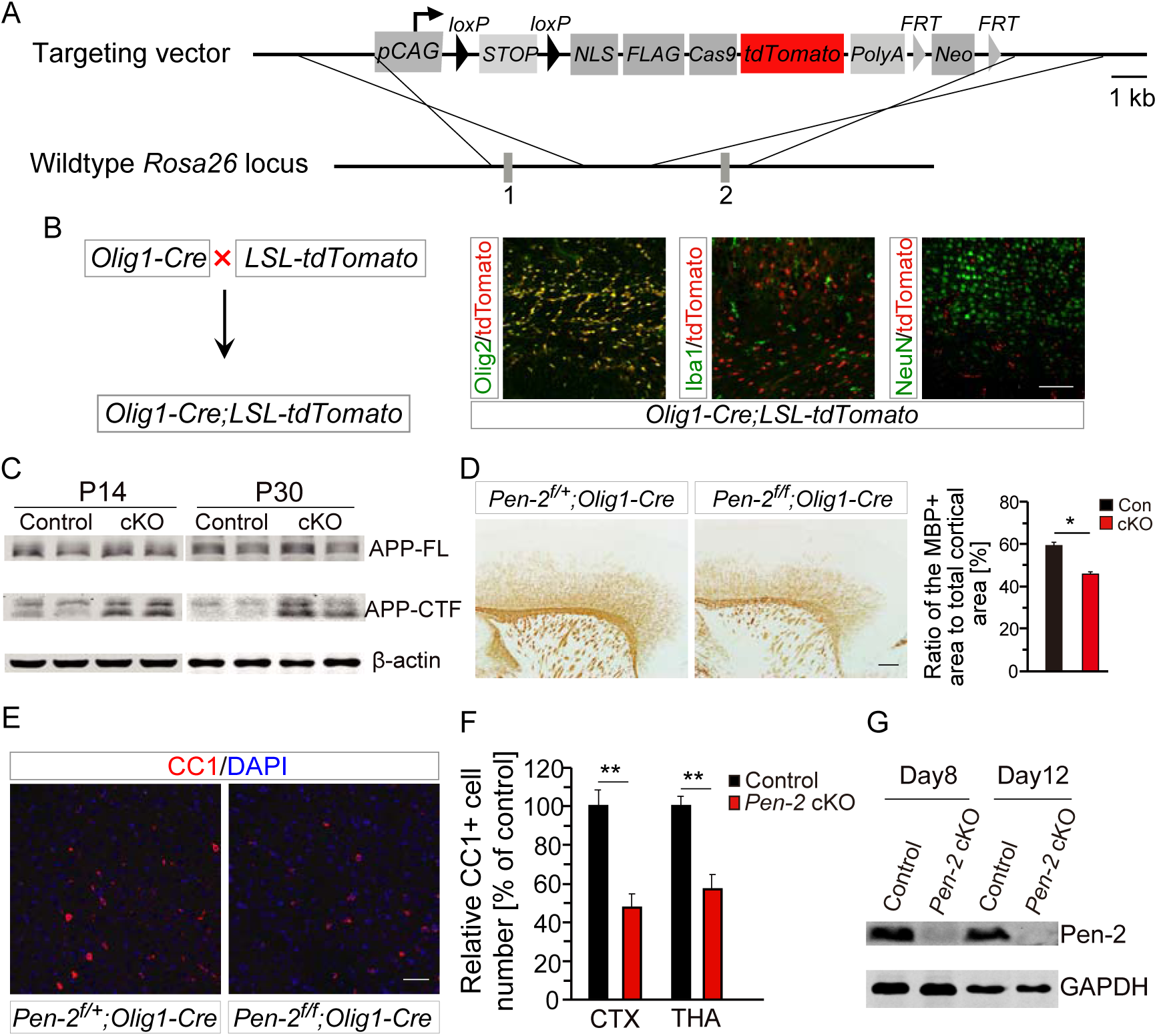
Loss of mature OLs in *Pen-2* cKO mice. A. Targeting strategy for generation of the tdTomato reporter mouse, which expresses tdTomato in a Cre-dependent manner. B. Characterization of Cre expression pattern using the tdTomato reporter mouse. Fluorescence IHC revealed that tdTomato is co-stained with Olig2 but not Iba1 or NeuN in the *Olig1-Cre* mouse expressing tdTomato at P2. C. Western blotting on APP-FL and APP-CTF using cortical samples. There was no significant difference on APP-FL levels between control and *Pen-2* cKO mice at either P14 or P30. There was highly significant difference on APP-CTF levels between control and *Pen-2* cKO mice. β-actin was used as the loading control (Control: n=3; *Pen-2* cKO: n=4). D. IHC on Mbp. There was significant difference on the ratio of the Mbp-positive area to the total cortical area between control and *Pen-2* cKO mice at P30 (Control: n=5; *Pen-2* cKO: n=6; p<0.05). E. Fluorescence IHC on CC1 in the cortex. Mice at P30 were used. F. Relative number of CC1+ cells in the cortex and the thalamus. There were significant differences on the averaged number of CC1+ cells in 1 mm^2^ area between control and *Pen-2* cKO mice at P30 (Control: n=4; *Pen-2* cKO: n=5; **, p<0.01). G. Western blotting using lysates prepared from OPC cultures. OPCs were cultured from control and *Pen-2* cKO cortices. Cell lysates were collected at OPC culture days 8 and 12. Pen-2 was not detected in *Pen-2* cKO cultures. Scale bar is 100 µm in (A) or 500 µm in (D) and (E).

**Figure EV2.**
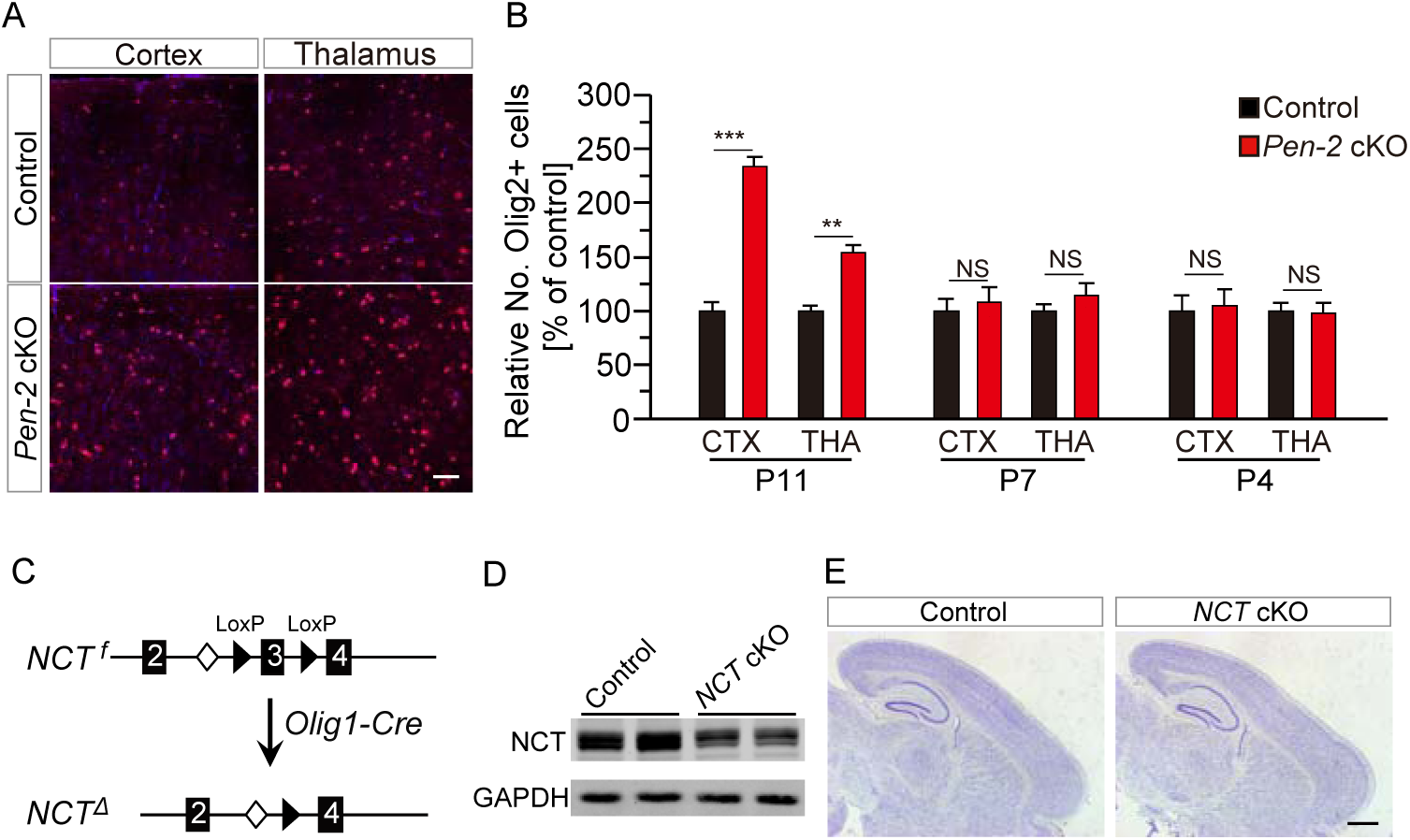
A. Age-related increase on the total number of OPCs in *Pen-2* cKO mice and the generation of *NCT* cKO mice. B. Representative images for Olig2 IHC in the cortex and the thalamus in control and *Pen-2* cKO mice at P11. C. Relative number of Olig2+ cells in the cortex and the thalamus in *Pen-2* cKO mice at various ages. There was significant difference on the averaged number of Olig2+ cells (in 1 mm^2^ area) between control and *Pen-2* cKO mice at P11 but not P4 and P7 (n=3 mice/group/age; **, p<0.01; ***, p<0.005; NS, not significant). D. Breeding plan for generation of OL lineages specific *NCT* cKO mice. E. Western blotting on NCT. There was significant difference on protein levels of NCT between control and *NCT* cKO mice (Control: n=3; *NCT* cKO: n=4). F. Nissl staining. There were no detectable changes in *NCT* cKO mice at P14 compared with controls. Scale bar is 50 µm in (A) or 1 mm in (E).

**Figure EV3.**
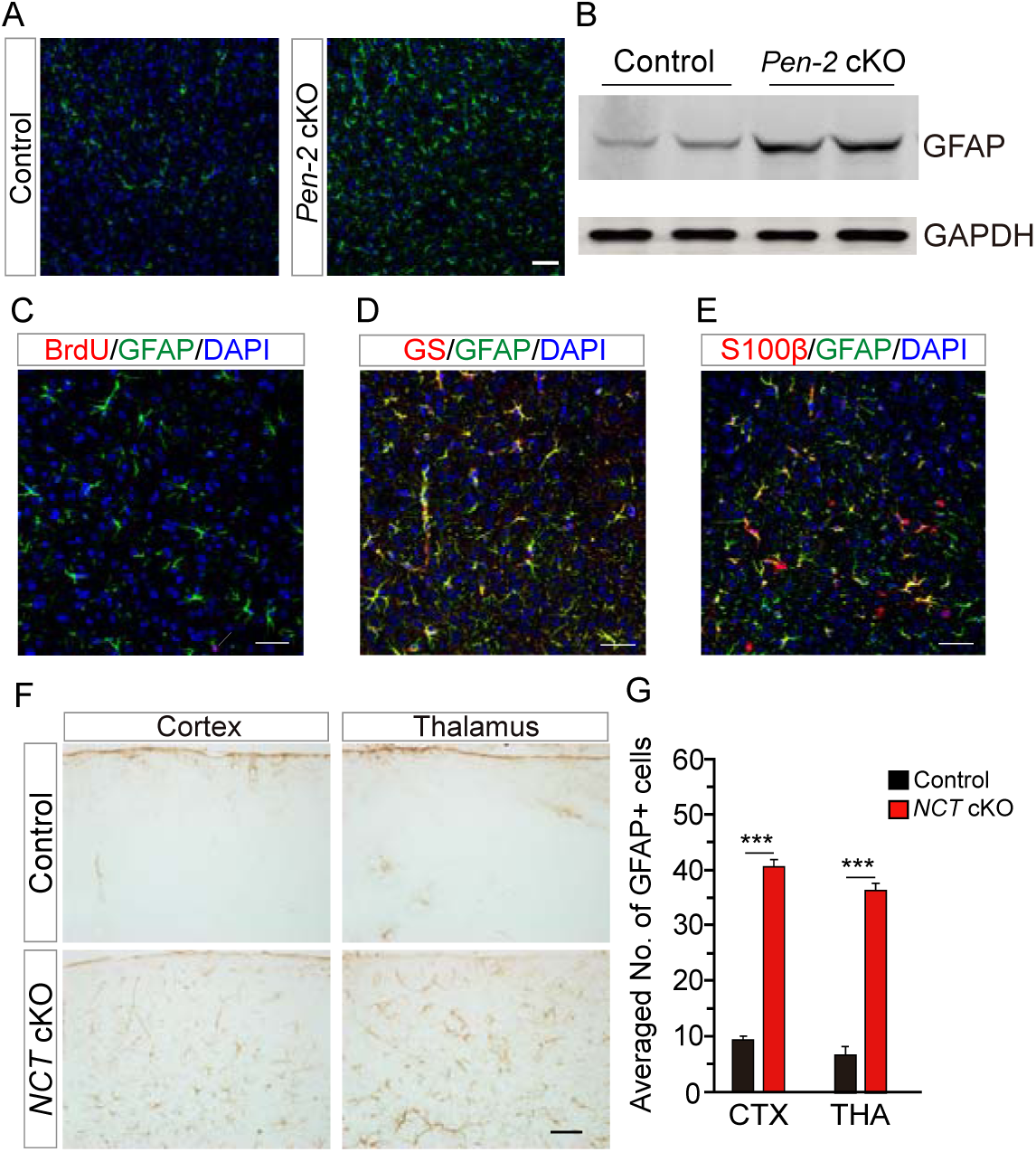
Enhanced astrogenesis in *Pen-2* cKO and *NCT* cKO mice. A. GFAP IHC in the spinal cord. There was increased immuno-reactivity of GFAP in *Pen-2* cKO mice at P14. B. Western blotting on GFAP. There were significantly increased levels of GFAP in the cortex of *Pen-2* cKO mice compared with controls at P14 (Control: n=3; *Pen-2* cKO: n=4). GAPDH served as the loading control. C. Double staining for GFAP and BrdU. BrdU was injected into mice at P14 for 1 hr. GFAP+/BrdU+ cells were not observed in cortices of control or *Pen-2* cKO mice. (D-E) Double staining for GFAP and GS (D) or S100β (E). GFAP+ cells were immuno-positive for GS (D) or S100β (E) in the cortex of *Pen-2* cKO mice. F. GFAP IHC in the cortex and the thalamus in *NCT* cKO mice at P14. G. Averaged number of GFAP+ cells in each microscopic field (under the 20×objective lens) in the cortex and the thalamus. There were significant differences between control and *NCT* cKO mice at P14 (n=3 per group; ***, p<0.005). Scale bar is 50 µm in (A), (C), (D) and (F).

**Figure EV4.**
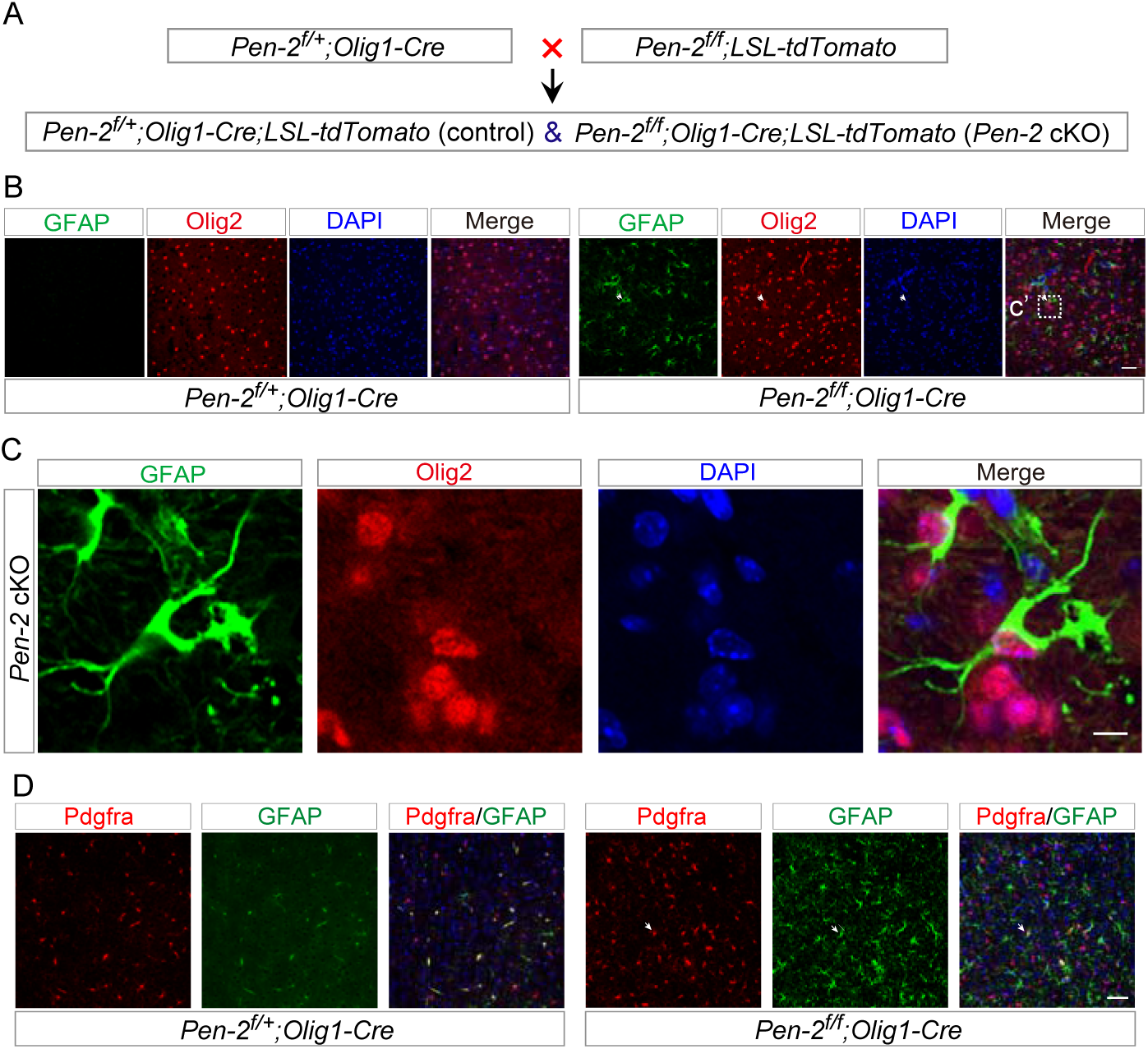
Generation of astrocytes from OPCs in *Pen-2* cKO mice. A. Breeding strategy for the generation of *Pen-2* cKO mice expressing tdTomato. *Pen-2^f/+^;Olig1-Cre;LSL-tdTomato* and *Pen-2^f/f^;Olig1-Cre;LSL-tdTomato* served as control and *Pen-2* cKO mice, respectively. B. Double staining for GFAP and Olig2 in the thalamus of control and *Pen-2* cKO mice at P14. Olig2 was co-stained with GFAP in *Pen-2* cKO but not control mice. C. The boxed area in (B) was enlarged here for co-localization for GFAP and Olig2 in *Pen-2* cKO mice. D. Double staining on Pdgfrα and GFAP. There were abundant Pdgfrα+/GFAP+ cells in the cortex of *Pen-2* cKO but not control mice at P12. Scale bar is 50 µm in (A) and (C), or 20 µm in (B).

**Figure EV5.**
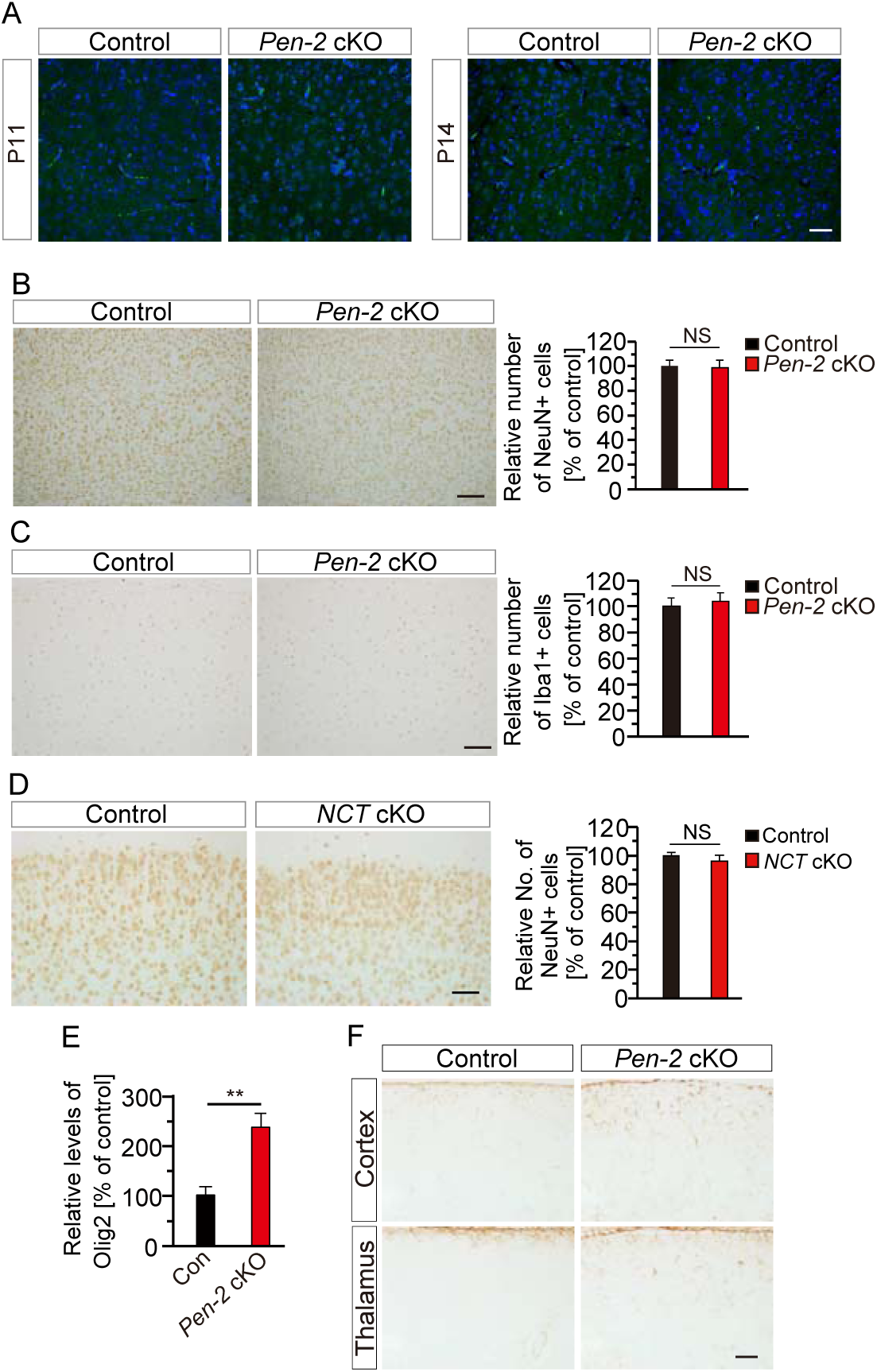
No neuronal loss in *Pen-2* cKO and *NCT* cKO mice. A. TUNEL staining for mice at P11 and P14. There were no TUNEL+ cells in the cortex or the thalamus in control and *Pen-2* cKO mice at each age. B. NeuN IHC. Relative number of NeuN+ cells (in 1 mm^2^ area) in the cortex of *Pen-2* cKO mice was not different from that in controls at P14. C. Iba1 IHC. There was no significant difference on relative number of Iba1+ cells (in 1 mm^2^ area) in the cortex between control and *Pen-2* cKO mice at P14 (n=4 per group; NS, not significant). D. NeuN IHC. There was no significant difference on relative number of NeuN+ cells (in 1 mm^2^ area) in the cortex between control and *NCT* cKO mice at P14 (n=3 per group; NS, not significant). E. Q-PCR analysis on *Olig2* mRNAs in cortical samples at P14. There was significant difference between control and *Pen-2* cKO cortices (n=4 per group; p<0.01). F. GFAP IHC in the mouse brain at P11. There was increased immuno-reactivity of GFAP in the cortex or the thalamus in *Pen-2* cKO mice compared with controls. Scale bar is 50 µm in (A), (B), (C), (D) and (F).

**Figure EV6.**
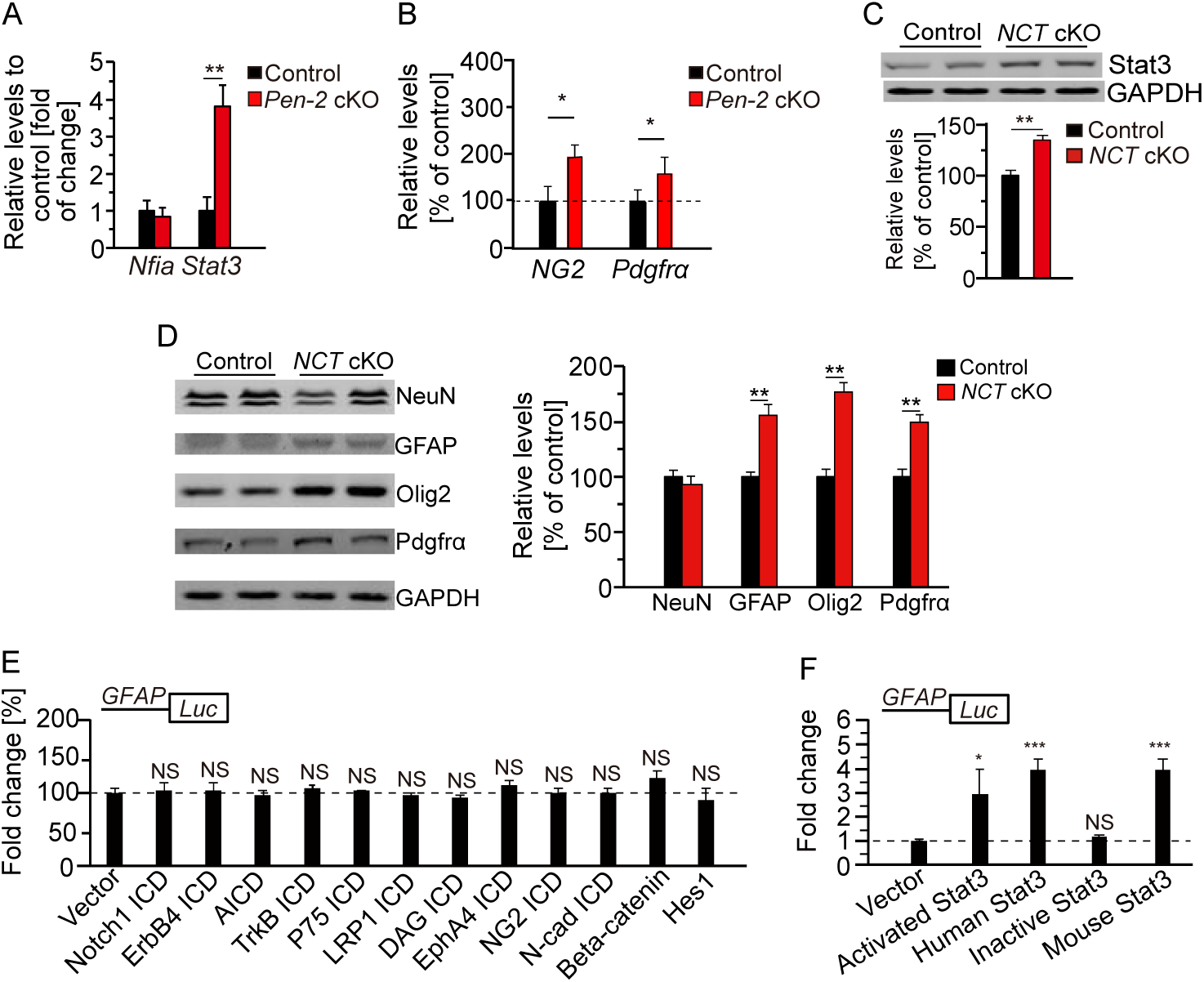
Increased expression of Stat3 and GFAP due to loss of γ–secrease function. A. Q-PCR analyses on *Nfia* and *Stat3* mRNAs in FACS-purified cells from cortices of *Pen-2^f/+^;Olig1-Cre;mTmG* and *Pen-2^f/f^;Olig1-Cre;mTmG* mice at P11. There was significant difference on *Stat3* but not *Nfia* between control and *Pen-2* cKO mice (Control: n=3 mice; *Pen-2* cKO: n=4; **, p<0.01). B. Q-PCR analyses on *NG2* and *Pdgfra* mRNAs in the cortex of control and *Pen-2* cKO mice at P14. There were significant differences between two groups (n=4 mice per group; *, p<0.05). C. Western analysis on Stat3 in *NCT* cKO mice at P14. There was significant difference between control and *NCT* cKO mice (n=4 per group; *, p<0.05). D. Western analyses on NeuN, GFAP, Olig2 and Pdgfra in *NCT* cKO mice at P14. There were significant differences on GFAP, Olig2 and Pdgfra but not NeuN between control and *NCT* cKO mice (n=4 per group; *, p<0.05). E. Luciferase assay on *GFAP* promoter activity. Expression of Notch1 ICD, APP ICD, TrkB ICD, P75 ICD, LRP1 ICD, DAG ICD, EphA4 ICD, NG2-ICD, N-cadherin ICD, β-catenin or Hes1 did not significantly affect the activity of the *GFAP* promoter (n=4 replicates; p>0.1). F. Luciferase assay on *GFAP* promoter activity. Expression of activated Stat3, human Stat3 and mouse Stat3 but not inactive Stat3 significantly inhibited the activity of the *GFAP* promoter (n=4 replicates; *, p<0.05; ***, p<0.005; NS, not significant).

**Figure EV7.**
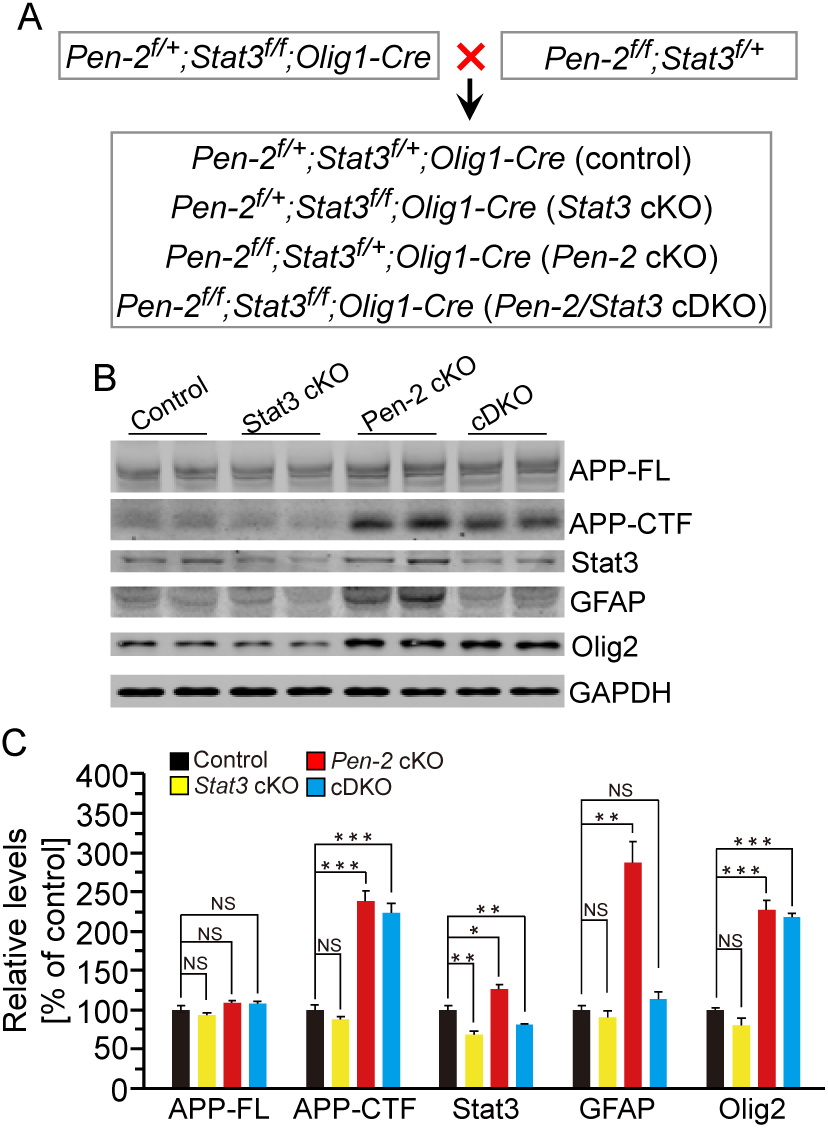
Restoration of astrocyte population in *Pen-2* cKO mice through OL lineages specific deletion of Stat3. A. Strategy for generation of *Pen-2/Stat3* cDKO mice. *Pen-2* cKO were bred with *Stat3^f/f^* to generate *Pen-2^f/+^;Stat3^f/+^;Olig1-Cre* and *Pen-2^f/+^;Stat3^f/+^*, which were inter-crossed to generate *Pen-2^f/+^;Stat3^f/f^;Olig1-Cre* and *Pen-2^f/f^;Stat3^f/+^*. The latter were crossed together to obtain control (*Pen-2^f/+^;Stat3^f/+^;Olig1-Cre*), *Stat3* cKO (*Pen-2^f/+^;Stat3^f/f^;Olig1-Cre*), *Pen-2* cKO (*Pen-2^f/f^;Stat3^f/+^;Olig1-Cre*) and *Pen-2/Stat3* cDKO (*Pen-2^f/f^;Stat3^f/f^;Olig1-Cre*) mice. B. Western analyses on APP-FL, APP-CTF, Stat3, GFAP and Olig2 in mice at P14. C. Quantification on APP-FL, APP-CTF, Stat3, GFAP and Olig2. APP-FL levels were not changed in *Stat3* cKO, *Pen-2* cKO and *Pen-2/Stat3* cDKO mice compared with controls at P14. There was no difference on APP-CTFs between *Pen-2* cKO and *Pen-2/Stat3* cDKO mice. Stat3 levels were reduced in *Stat3* cKO and *Pen-2/Stat3* cDKO compared with control and *Pen-2* cKO. GFAP levels were reduced in *Pen-2/Stat3* cDKO compared with *Pen-2* cKO. Olig2 levels were not reduced in *Pen-2/Stat3* cDKO compared with *Pen-2* cKO. GAPDH was used as the loading control (Control: n=3; *Stat3* cKO: n=3; *Pen-2* cKO: n=3; *Pen-2/Stat3* cDKO: n=4; *, p<0.05; **, p<0.01; ***, p<0.005; NS, not significant).

